# Cigarette smoke aggravates atherosclerosis by promoting the infiltration of inflammasome-primed neutrophils and disrupting macrophage function in lesions

**DOI:** 10.1101/2025.10.14.682475

**Authors:** Dipanjan Chattopadhyay, Nitin Nitin, Robert M. Jaggers, Baskaran Athmanathan, Krishna P Maremanda, Babunageswararao Kanuri, Albert Dahdah, Rajnikant Shukla, Man SK Lee, Andrew J. Murphy, Prabhakara R. Nagareddy

**Author notes:** **Corresponding author,** Dr. Prabha Nagareddy, PhD, FAHA Department of Medicine, Cardiovascular Section University of Oklahoma Health Sciences Center, Oklahoma City, USA Ex 44737. These authors contributed equally (Dipanjan Chattopadhyay, Nitin Nitin, Robert M. Jaggers, Baskaran Athmanathan).

## Abstract

**Background:** Cigarette smoking (CS) is a major risk factor for cardiovascular disease (CVD) through chronic inflammation. While its pulmonary effects are well established, the mechanisms linking lung inflammation to vascular injury remain unclear. Because neutrophils are early responders to CS-induced inflammation, we hypothesized that they drive systemic myelopoiesis and vascular inflammation via alarmin release.

**Methods:** Wild-type (WT) mice were exposed to inhaled CS or orally administered cigarette smoke extract (CSE). Immune cell composition in lung, bronchoalveolar lavage fluid (BALF), blood, spleen, and bone marrow (BM) was assessed by flow cytometry. Hematopoietic stem and progenitor cell (HSPC) proliferation, reactive oxygen species (ROS) production, and S100A8/A9 release were quantified. Atherosclerosis progression was evaluated in *Ldlr*^⁻^*^/^*^⁻^ mice fed a Western diet and treated with CSE. To define the role of neutrophil-derived S100A8/A9, bone marrow transplantation was performed using *S100a9*^⁻^*^/^*^⁻^ or WT donors.

**Results:** CS exposure increased circulating monocytes and neutrophils through enhanced BM myelopoiesis and elevated ROS-dependent S100A8/A9 release. Oral CSE reproduced these effects, indicating direct activation of neutrophils independent of pulmonary inflammation or lipid changes. In *Ldlr*^⁻^*^/^*^⁻^ mice, CSE accelerated atherosclerosis by promoting infiltration of inflammasome-primed neutrophils, increased IL-1β release, and impaired macrophage efferocytosis. Hematopoietic *S100a9* deletion normalized myelopoiesis and reduced vascular inflammation and plaque burden.

**Conclusions:** Ingested CS components directly activate neutrophils to release S100A8/A9, triggering myelopoiesis and vascular inflammation. These findings reveal that tobacco’s cardiovascular toxicity extends beyond inhalation, implicating oral exposure as a driver of systemic inflammation and atherogenesis.

**Novelty and Significance:** *What Is Known?:*

- Cigarette smoking (CS) is a major risk factor for atherosclerosis, driving systemic inflammation and innate immune activation.
- Neutrophils and monocytes contribute to plaque progression, but the upstream mechanisms by which CS exacerbates their pathogenic roles remain incompletely understood.
- S100A8/A9 levels correlate with neutrophilia and cardiovascular risk in smokers, but their functional role in lesion biology is not fully defined.

*What New Information Does This Article Contribute?:*

- Identifies S100A8/A9 as a key mediator linking CS exposure to enhanced medullary myelopoiesis, neutrophilia, and increased lesional infiltration of inflammasome-primed myeloid cells.
- Demonstrates that neutrophil-derived IL-1β impairs macrophage efferocytosis by downregulating phagocytosis receptors, thereby promoting plaque vulnerability.
- Reveals that CS drives atherosclerosis even in the absence of lipid perturbations or overt pulmonary injury, highlighting a novel oral exposure–vascular axis of disease propagation.

## Introduction

The association of cigarette smoking (CS) and cardiovascular disease (CVD) is well-known. The risk of acute coronary and cerebrovascular events, including myocardial infarction (MI), stroke and sudden death is markedly increased by smoking (CS)^1,2^. This is likely due to indiscriminate and accelerated atherogenesis in various arteries including the coronary, carotid, cerebral, and peripheral vasculature^3^. Although smoking is responsible for nearly 10% of all CVD cases^4^, the precise cellar and molecular events that link CS with atherosclerosis is not clear. Cigarette smoke contains more than 4000 chemicals and particulate matter that makes it hard to identify the exact factor (s) responsible for CVD risk^5^. Nonetheless, over the years many mechanisms have been postulated to describe mechanistic links between CS and atherosclerosis disease burden. These include direct deleterious effects on cells in the vascular wall, the components of blood including lipoproteins and immune cells. For example, CS smoke is associated with elevated neutrophil and monocyte counts^6^ likely due enhanced myeloid cell output from the bone marrow (BM)^7^. It is also known to induce direct structural and functional abnormalities in flow-mediated vasodilatation, an early marker of vascular and endothelial dysfunction^8,9^. CS is associated with alterations in serum lipid profiles including decreased HDL and increased VLDL, LDL and TG levels, all of which are known to promote atherogenesis ^10,11,12^. In addition, CS can also directly induce lipid oxidation resulting in the formation of oxidized LDL that is readily scavenged by macrophages and transform into foam cells during atheroslerosis^13^. Further, CS increases the expression of adhesion molecules on endothelial cells that help recruit inflammatory leucocytes^14^ further aiding and abetting atherosclerosis^15–17^. These derangements ranging from increased myeloid cell number to endothelial dysfunction and abnormal serum lipids all work collectively to enhance inflammation in the vessel wall and thus contribute to increased atherosclerosis.

Aside from the direct effect of CS on individual components of the CV system, it has much more deleterious impact on several cell types within the lungs including epithelial, alveoli, and immune cells^5^. Airway epithelial cells act as a barrier to invading pathogens and secretes mucus that helps to trap invading pathogens and inhaled particles^18^. This is achieved via secretion of various antimicrobial peptides^19^, chemokines and cytokines that invariably act as triggers of inflammation in the lungs^20,21^. Upon initiation of the inflammatory response, neutrophils are the first responders to the insult where they accumulate in large numbers and secrete reactive oxygen species (ROS), matrix metalloproteinases, and other enzymes. Although these products are beneficial for elimination of the invading dangers, they are also toxic and aid in alveolar destruction^22,23^. Amongst the cytokines produced during the initial phase of lung inflammation are TNF-α and IL-1β, that play a key role in the initiation of the inflammatory response on site but also have wide ranging systemic effects on the immune system^24,25^. In rodents, CS and lung inflammation are shown to be correlated and dependent on IL-1β signaling. Blocking of the NLRP3 inflammasome signaling that generate IL-1β in rodents subjected to CS reduced the inflammation and the neutrophilia^26^. Taken together these findings suggest a mechanistic link between local inflammation that occur within the lungs and system-wide inflammation as a result of increased immune cell output, which is a major driver of CVD.

Previous studies from our lab and elsewhere have shown that neutrophil-borne alarmins such as S100A8/A9 interact with their receptors on hematopoietic stem and progenitor cels (HSPCs) in the BM to promote myelopoiesis^27^. Because, the potential for recruitment of immune cells into plaque is strongly influenced by their numbers in circulation^28–31^, we reasoned that enhanced leukocytosis induced by CS-exposed neutrophils in the lung or blood would lead to greater atherosclerosis burden. Whether inflammation that occurs locally within the lung or systemically is the causative factor in CS-induced myelopoiesis is unknown. Therefore, we sought to investigate the signaling pathways that contribute to enhanced myelopoiesis in response to CS and studied its impact on atherosclerosis.

## Methods

Detailed methods including statistics are provided in the online-only Data Supplement^32^. All animal studies were approved by the IACUCs in accordance with the NIH Guide for the Care and Use of Laboratory Animals. Detailed analytical methods and reagents will be made available to other investigators on a reasonable request. Key resources and reagents used in the study are listed in Supplemental Table I (*Online-only Data Supplement*).

## Results

### Exposure to cigarette smoke (CS) in healthy wild-type (WT) mice induces leukocytosis and triggers the release of the alarmins S100A8/A9

To assess the effect of inhaled cigarette smoke on leukocyte levels, we conducted a temporal study in healthy mice exposed to either air or CS for 4 weeks. Exposure to CS led to a significant increase in circulating monocytes primarily the proinflammatory Ly6C^hi^ subset and neutrophils starting from week 2 (**Figures 1B through 1E**). Furthermore, neutrophils from CS-exposed mice exhibited increased surface expression of CD11b, a marker of cellular activation enhanced adhesive and migratory capacity (**Figure 1F**). The increase in number of leukocytes was accompanied by a significant rise in serum levels of S100A8/A9, an alarmin complex highly expressed in neutrophils (**Figure 1G**). To determine whether neutrophils are the primary source of circulating S100A8/A9, we assessed both plasma and intracellular S100A9 levels in blood neutrophils by flow cytometry (**Figure 1H and 1I**). We compared neutrophil S100A9 mean fluorescence intensity (MFI) between vehicle-treated and CS-exposed mice. The air-exposed group showed significantly higher neutrophil S100A9 MFI (35.31 ± 5.26) compared to the smoking group (21.97 ± 6.35), as determined by unpaired t-test (p = 0.0029) and confirmed by Mann-Whitney U test (p = 0.0087). To assess the relationship between plasma S100A8/A9 levels and neutrophil S100A9 MFI, we performed correlation analyses (**Figure 1J**). Across all samples, there was a strong and significant negative correlation (Spearman ρ = −0.73, p = 0.0065), indicating that higher plasma S100A8/A9 levels were associated with lower neutrophil S100A9 MFI. These findings suggest that CS exposure reduces neutrophil S100A9 expression and supports the conclusion that neutrophils are the major contributors to elevated S100A8/A9 levels in CS-exposed mice.

**Figure 1:**
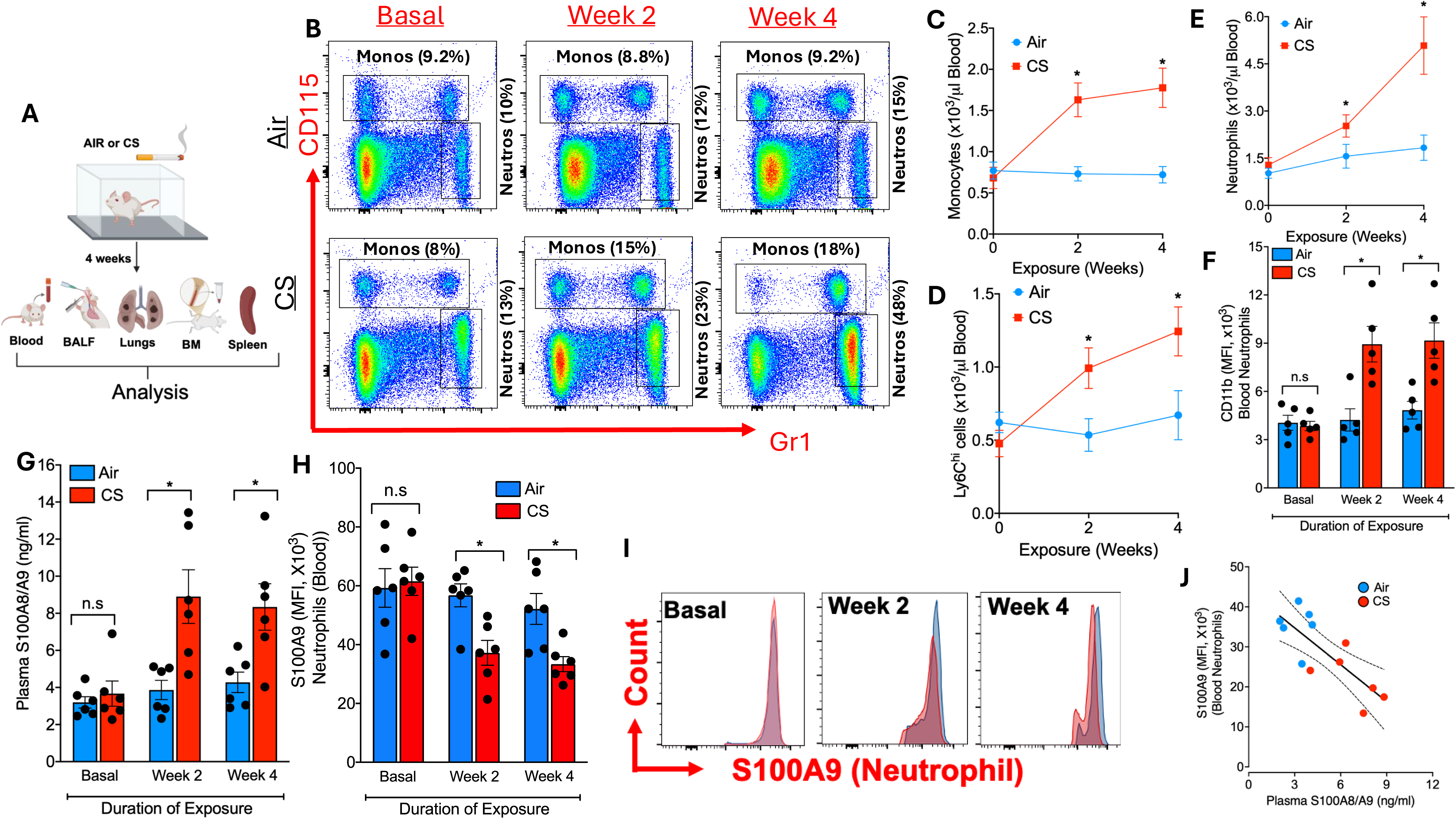
Cigarette smoke (CS) exposure induces leukocytosis and promotes S100A8/A9 release from activated neutrophils. **A**, Schematic overview of the experimental protocol: wild-type (WT) mice were exposed to either room air or cigarette smoke (CS) for up to 4 weeks. **B**, Representative flow cytometry plots illustrating the gating strategy for circulating leukocyte subsets. Quantification of total monocytes (**C**), proinflammatory Ly6C^hi^ monocytes (**D**), and neutrophils (**E**) over time in response to CS exposure. **F**, Mean fluorescence intensity (MFI) of CD11b expression on blood neutrophils, assessed by flow cytometry, indicating enhanced activation. **G**, Plasma concentrations of the neutrophil-derived alarmin complex S100A8/A9. Quantification (**H**) and representative histograms (**I**) of intracellular S100A9 levels in blood neutrophils, measured by flow cytometry. **J**, Correlation analysis between plasma S100A8/A9 levels and neutrophil S100A9 MFI. A linear regression line is included for visualization; statistical significance was assessed using Spearman’s rank correlation. Data represent mean ± SEM. Statistical analysis for **C–E**, Two-way repeated measures ANOVA with Bonferroni’s post hoc test (*P < 0.05 vs. air-exposed group at matched time points). **F–H**, Unpaired two-tailed t-test (*P < 0.05 vs. air; n.s., not significant). **J**, Spearman correlation revealed a significant inverse relationship between plasma S100A8/A9 and neutrophil S100A9 MFI (ρ = −0.73, *P* = 0.0065), suggesting neutrophils are a major source of extracellular S100A8/A9 following CS exposure.

### Leukocytosis in cigarette smoke (CS) exposed mice is due to enhanced myelopoiesis

Cigarette smoke skews neutrophil development at the progenitor level (early GMPs) in BM, with major transcriptional reprogramming. CYTOF profiling of BM progenitor cells show that GMPs are transcriptionally rewired to overproduce pro-inflammatory neutrophils^33^. To understand the cellular basis for the increased circulating monocytes and neutrophils observed in our CS-exposed mice, we assessed the proliferation of HSPCs including GMPs in the BM. This was performed by administering EdU, a thymidine analog incorporated during active DNA synthesis, shortly before euthanasia. As expected, CS-exposed mice exhibited a marked increase in HSPC proliferation, particularly within the common myeloid progenitors (CMPs) and granulocyte-macrophage progenitors (GMPs), which are key precursors of monocytes and neutrophils (**Figure 2A and 2B**). These results were further validated using additional proliferation markers, including *Ki-67, Pu.1*, and *Ptprc* (**Figure 2C**). We then compared the numbers of monocytes and neutrophils in the BM of CS- and air-exposed mice. Although the total numbers of these cells were similar between the two groups (**Figure 2D and 2E**), the number of newly generated monocytes and neutrophils was significantly higher in the CS group (**Figure 2D and 2F**). We next examined spleen as a potential source of extramedullary myelopoiesis. Interestingly, the number of HSPCs (**Figure S1A**) or their proliferation (**Figure S1B**) in spleen were unaffected. Similarly, the number of monocytes and neutrophils (**Figure S1C**) did not change. However, the prolonged exposure to CS led to a significant decrease in splenic weight (**Figure S1D**), a likely indication of immunosuppression and/ or redistribution of immune cells (e.g., neutrophilia in blood but reduced splenic cellularity). Intriguingly, intracellular levels of S100A8/A9 were elevated in splenic neutrophils (**Figure S1E**), suggesting that the spleen may serve as an additional site of alarmin production or storage in response to CS exposure. These findings suggest that CS-induced leukocytosis is primarily driven by increased myelopoiesis in the BM and not spleen.

**Figure 2:**
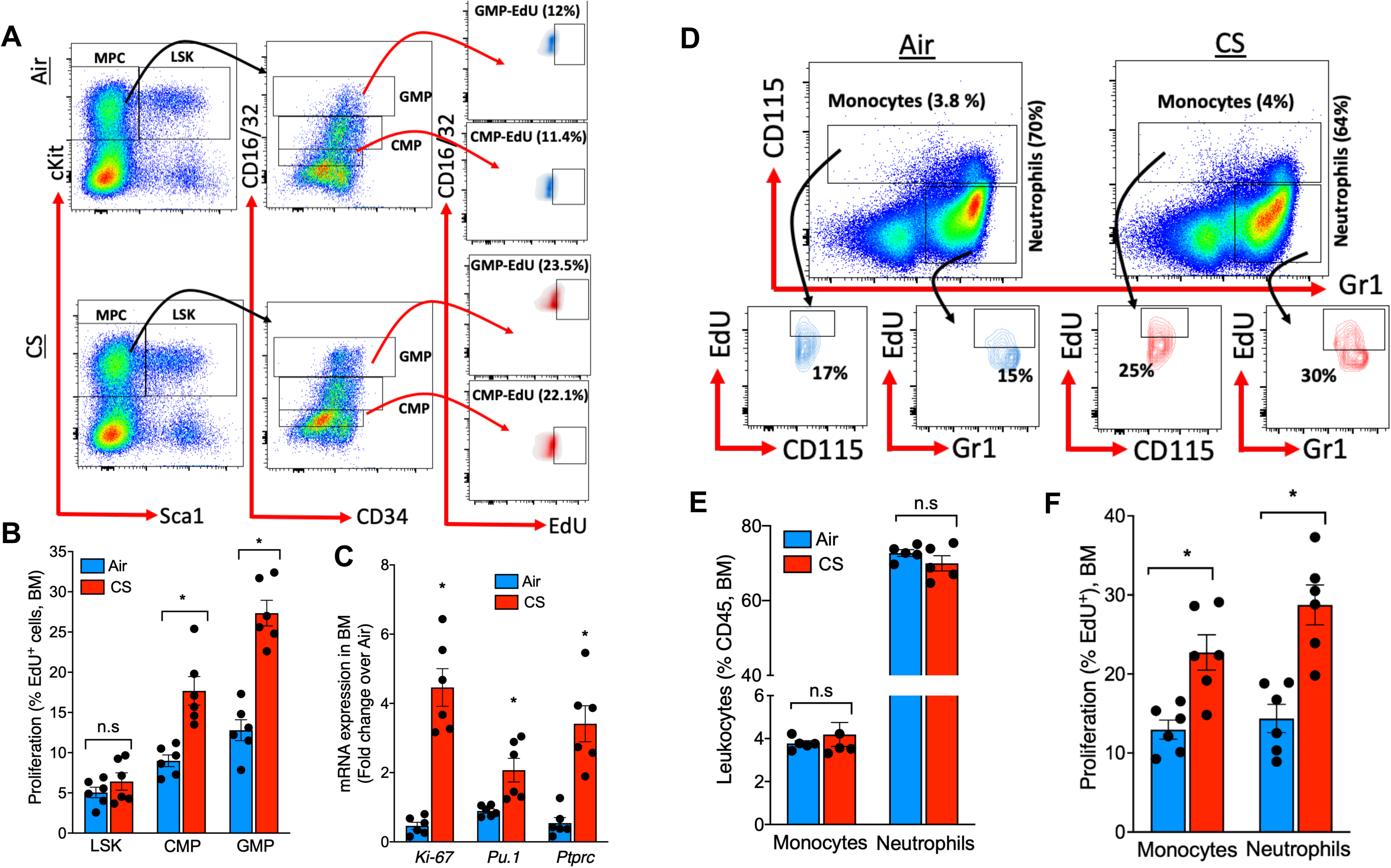
Cigarette smoke (CS) stimulates myelopoiesis, leading to elevated number of leukocytes counts. **A)**, Representative flow cytometry plots showing the gating strategy used to identify and quantify myeloid progenitor populations in the bone marrow, including Lineage⁻ C-Kit⁺ Sca1⁻ (LSK) cells, common myeloid progenitors (CMPs), and granulocyte-macrophage progenitors (GMPs). Mice were injected with EdU 12 hours before sacrifice to assess proliferation; values in parentheses indicate the percentage of EdU⁺ cells. **B)**, Quantification of proliferating (EdU⁺) LSKs, CMPs, and GMPs. **C)**, Gene expression analysis of key myelo-proliferative markers (*Ki-67*, *Pu.1*, and *Ptprc*) in BM cells, assessed by RT-PCR. **D)**, Representative flow cytometry plots showing gating strategies for identifying monocytes and neutrophils in the BM, with EdU administered 12 hours prior to euthanasia. **E)**, Quantification of total monocytes and neutrophils in the BM, and **F)**, percentage of each population that incorporated EdU. All data were gated to exclude debris, dead cells, and doublets. Results are expressed as mean ± SEM. Statistical analysis for B -F, Unpaired two-tailed t-test (*P < 0.05 vs. corresponding air-exposed group; n.s., not significant).

### Cigarette smoke (CS) exposure led to significant increase in lung inflammation

To explore how inhaled CS promotes BM myelopoiesis and the release of myeloproliferative cytokines such as S100A8/A9, we assessed markers of inflammation in the lungs of air- and CS-exposed mice. Both bronchoalveolar lavage fluid (BALF) and lung tissue were analyzed at the study endpoint. CS exposure led to a marked increase in neutrophils, interstitial macrophages (IMs), CD103⁺ dendritic cells (DCs), and Ly6C⁺ monocytes in lung tissue, while the numbers of alveolar macrophages (AMs), CD11b⁺ DCs, eosinophils, and Ly6C⁻ monocytes were reduced (**Figure S2A through S2C**). A similar proinflammatory pattern was observed in the BALF, with elevated levels of total leukocytes, neutrophils, CD103⁺ DCs, IMs, and Ly6C⁺ monocytes (**Figure S2D**). However, in contrast to lung tissue, the numbers of AMs and Ly6C⁻ monocytes in the BALF remained unchanged. These findings indicate that CS exposure drives a proinflammatory shift in the lung immune environment, characterized by increased recruitment of inflammatory myeloid cells and a reduction in homeostatic, anti-inflammatory populations. In support of this, we also observed elevated levels of S100A8/A9 in the BALF and increased expression of its receptors, TLR4 and RAGE, in lung tissue (**Figure S2E and S2F**). Together, these results suggest that CS triggers inflammatory remodeling of the lung, which may serve as a source of systemic inflammatory cues particularly S100A8/A9 that stimulate BM activation and promote myelopoiesis.

### Circulating inflammatory cues generated during CS exposure drive enhanced myelopoiesis in the BM

To determine whether the inflammatory cues activating HSPCs in the BM originate from the lung or the systemic circulation, we conducted an oral gavage study in which healthy WT mice were administered two doses of cigarette smoke extract (CSE) over the same duration as the inhalation exposure. Mice were treated with either vehicle, 0.4 mg/kg, or 8 mg/kg of CSE for 4 weeks, during which leukocyte counts were monitored to assess the impact on leukocytosis and myelopoiesis (Experimental outline, **Figure 3A**). Oral administration of CSE resulted in a dose-dependent increase in circulating monocytes (**Figure 3B and 3C**) and neutrophils (**Figure 3B and 3D**). Consistent with findings from CS-inhalation studies, CSE treatment also led to a significant rise in plasma levels of S100A8/A9 (**Figure 3E**), likely due to secretion by circulating neutrophils. Neutrophil S100A9 MFI decreased significantly in a dose-dependent manner in CSE-treated groups (**Figure 3F**). Plasma S100A8/A9 levels showed a strong inverse correlation with neutrophil S100A9 MFI (Spearman ρ = −0.66, p = 0.0006), indicating that CSE exposure reduces neutrophil S100A9 expression and that plasma S100A8/A9 acts as a negative predictor of neutrophil S100A9 MFI (**Figure 3G**).

**Figure 3:**
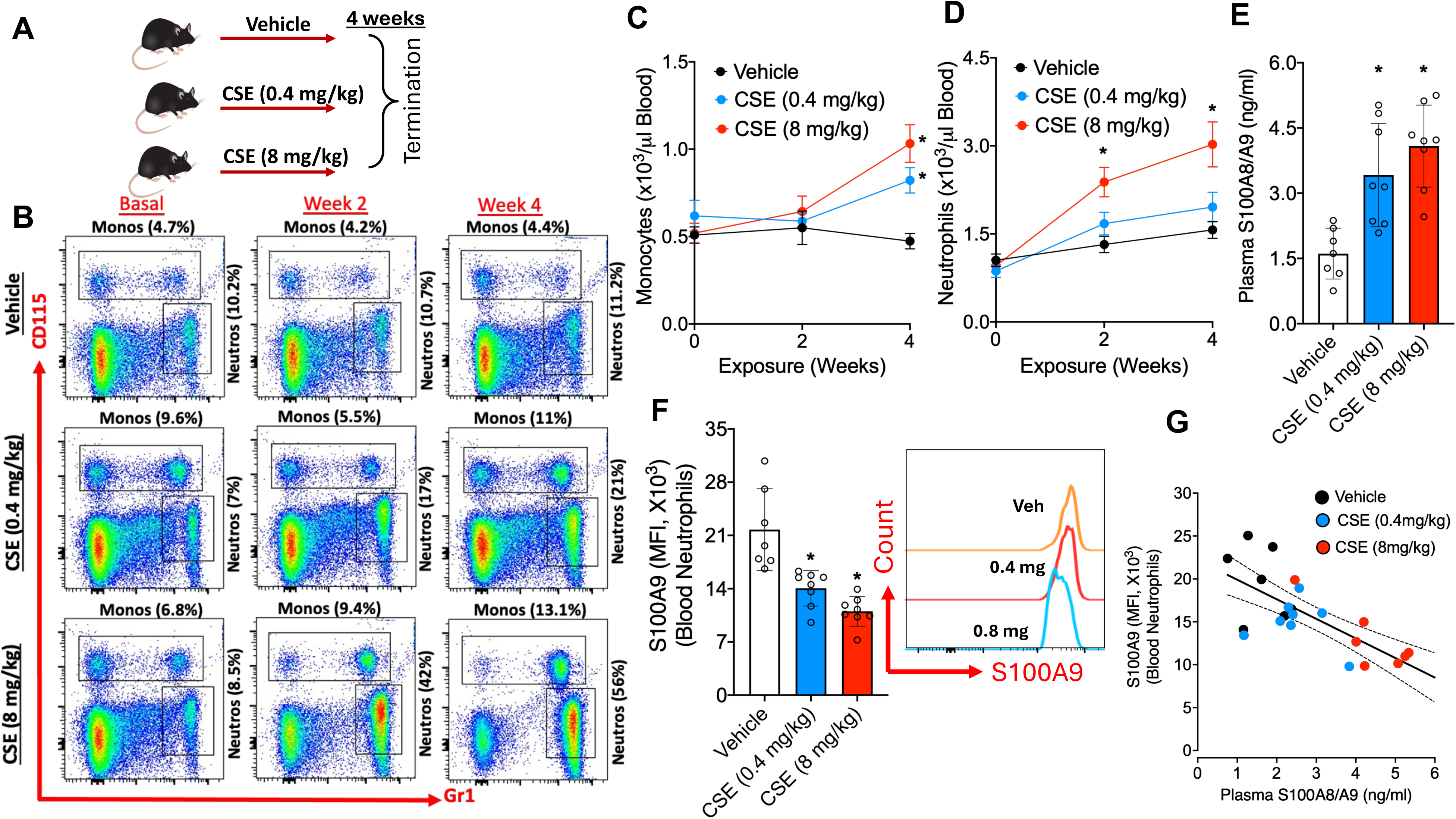
Circulating inflammatory cues generated during CS exposure drive enhanced myelopoiesis in the BM. **A)**, Schematic of the experimental protocol: Wild-type (WT) mice were orally gavaged with vehicle, low-dose CSE (0.4 mg/kg), or high-dose CSE (8 mg/kg) twice daily for 4 weeks. **B)**, Representative flow cytometric plots showing blood leukocyte populations in vehicle- and CSE-treated groups. Quantification of circulating total monocytes (**C**) and neutrophils (**D**) at different time points during CSE gavage. **E)**, Plasma levels of S100A8/A9 as measured by ELISA. **F)**, Mean fluorescence intensity (MFI) of intracellular S100A9 in neutrophils, as measured by flow and illustrated by histograms. **G)**, Spearman correlation analysis showing a strong inverse relationship between plasma S100A8/A9 levels and neutrophil S100A9 MFI, indicating that as neutrophils release S100A8/A9, intracellular stores are depleted. A linear regression line is shown for visualization. Data are presented as mean ± SEM. Statistical analysis for C and D (Two-way repeated measures ANOVA with Bonferroni’s post hoc test), E and F (one-way ANOVA with Tukey’s multiple comparison test (*p < 0.05, vs. vehicle). For G, Spearman’s rank correlation coefficient was used (ρ = −0.66, p = 0.0006).

To determine whether elevated blood leukocyte counts in CSE-treated mice were driven by increased BM myelopoiesis, we analyzed progenitor cell proliferation. As anticipated, oral CSE administration resulted in a dose-dependent increase in the proliferation of CMPs and GMPs in the BM (**Figure S3A and S3B**). This was further supported by upregulation of proliferation and myeloid lineage markers, including *Ki67, Pu.1, and Ptprc* (**Figure S3C**). Enhanced myelopoiesis was also evident from the increased production of myeloid cells (**Figure S3E**), particularly proinflammatory Ly6C^hi^ monocytes (**Figure S3F**) and immature neutrophils with elevated CD62L expression (**Figure S3G**). As expected, oral CSE exposure did not impact lung mass (**Figure S4A**), total lung cellularity (**Figure S4B**), or the overall composition of immune cells within lung tissue (**Figure S4C**). However, despite being given orally, CSE induced mild signs of lung inflammation (**Figures S4D and S4E**), likely due to the infiltration of activated neutrophils from the circulation (**Figure S4F**). This was further supported by the observation that neutrophils from both the blood and lung showed similar expression levels of inflammatory markers (**Figure S4G**). Together, these findings suggest that certain components of cigarette smoke whether inhaled or ingested generate systemic inflammatory cues that act on HSPCs in the BM to promote myelopoiesis.

### Interaction of neutrophil-derived S100A8/A9 with its putative receptors on HSPCs in the BM promote myelopoiesis

Previous studies from our lab and others have shown that neutrophils under stress can release the alarmin complex, S100A8/A9^27,34^. Given the significant elevation of plasma S100A8/A9 levels observed in both CS-exposed and CSE-treated mice, we hypothesized that CSE directly induces its release from neutrophils. To test this, we treated BM–neutrophils with increasing concentrations of CSE and observed a robust, dose-dependent increase in S100A8/A9 secretion (**Figure S5A**). Since oxidative stress is known to trigger S100A8/A9 release, we compared the effects of CSE and phorbol myristate acetate (PMA), a known inducer of S100A8/A9 in the presence or absence of diphenyleneiodonium (DPI), an NADPH oxidase inhibitor^34^. Both CSE and PMA induced strong S100A8/A9 release, which was significantly suppressed by DPI pretreatment (**Figure S5B)**. These findings suggest that CSE induces oxidative stress in neutrophils, leading to the release of S100A8/A9. This proinflammatory signal likely contributes to enhanced myelopoiesis in the BM. To confirm this mechanism *in vivo*, we measured reactive oxygen species (ROS) levels in neutrophils from various tissues in both CS- and CSE-treated mice. As expected, ROS levels in BM and splenic neutrophils were significantly lower than in circulating neutrophils in both air-exposed (**Figure S5C and S5D**) and vehicle-treated mice (**Figure S5E**). Notably, exposure to CS (by inhalation) or CSE (via gavage) significantly increased ROS levels in blood neutrophils, while ROS levels in BM and splenic neutrophils remained unchanged. These results suggest that systemic oxidative stress, particularly in circulating neutrophils, may underlie the release of S100A8/A9 in response to cigarette smoke exposure.

We next investigated whether neutrophil-derived S100A8/A9 alone is sufficient or if CS exposure is required to stimulate myelopoiesis. To test this, we cultured freshly isolated BM cells from WT-mice in the presence of various growth factors and treated them with either S100A8/A9 (2 µg/ml), CSE (20 µg/ml), or both (Experimental outline, **Figure 4A**). Cell proliferation was assessed by measuring EdU incorporation in different HSPC populations using flow cytometry. As shown previously^35^, stimulation of BM cells with S100A8/A9 alone had no effect on HSPC proliferation. Surprisingly, treatment with CSE alone also failed to enhance myelopoiesis. However, when both S100A8/A9 and CSE were combined, there was a significant increase in the proliferation of CMPs and GMPs (**Figure 4B and 4C**). Given that the primary effect of CS is to induce oxidative stress in neutrophils and thereby promote the release of S100A8/A9, we hypothesized that CS may also influence the expression of DAMP receptors on BM cells. To test this, we assessed the surface expression of TLR4 and RAGE, the two main receptors for S100A8/A9 and observed a marked increase in TLR4 expression, particularly on CMPs and GMPs (**Figure 4D**). These findings suggest that CS not only stimulates ROS production in neutrophils to trigger S100A8/A9 release but also upregulates TLR4 expression on progenitor cells (**Figure 4E**), thereby enhancing S100A8/A9-TLR4 interactions that drive myelopoiesis. We confirmed the functional role of S100A8/A9-TLR4 signaling by showing that BM cells from *Tlr4*^⁻^*^/^*^⁻^ mice failed to exhibit increased myelopoiesis in response to CSE+ S100A8/A9 treatment (**Figure 4F and 4G**).

**Figure 4:**
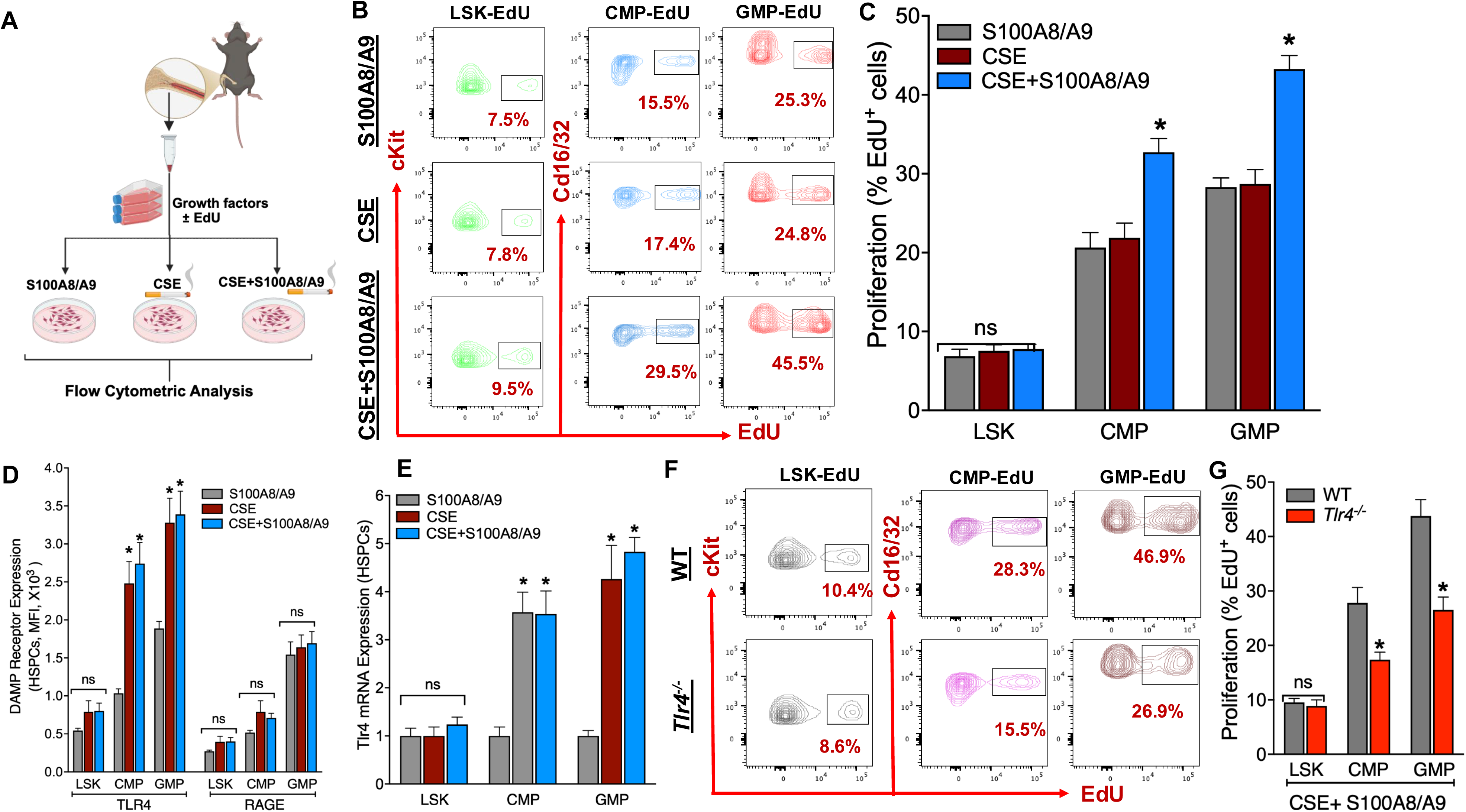
Synergistic activation of myelopoiesis by S100A8/A9 and CSE is mediated via TLR4 upregulation on myeloid progenitors. **A)**, Schematic of experimental design: Wild-type (WT) bone marrow (BM) cells were cultured with hematopoietic growth factors and treated with recombinant S100A8/A9 (2 µg/ml), cigarette smoke extract (CSE, 20 µg/ml), or both for 24 hours. Proliferation of hematopoietic stem and progenitor cells (HSPCs) and expression of damage-associated molecular pattern (DAMP) receptors were assessed by PCR and flow cytometry. **B)**, Representative flow cytometry plots showing EdU incorporation in common myeloid progenitors (CMPs) and granulocyte-macrophage progenitors (GMPs) following treatment. **C)**, Quantification reveals that only the combination of S100A8/A9 and CSE significantly enhances proliferation of CMPs and GMPs, while either agent alone has minimal effect. **D)**, Mean fluorescence intensity (MFI) of surface TLR4 and RAGE expression on CMPs and GMPs demonstrates selective upregulation of TLR4 in response to CSE. **E)**, Tlr4 mRNA expression across HSPC subsets confirms increased transcriptional induction of TLR4 by CSE. **F)**, Representative flow plots depicting EdU incorporation in WT and *Tlr4*^⁻^*^/^*^⁻^ HSPCs (LSKs, CMPs, and GMPs) treated with S100A8/A9, CSE, or both. **G)**, Quantification of EdU⁺ cells across HSPC subsets shows that only WT BM cells respond to S100A8/A9 + CSE co-treatment with enhanced myelopoiesis, confirming a critical role for TLR4 in mediating progenitor proliferation. Data represent mean ± SEM. Statistical analysis: **C–E**, one-way ANOVA with Tukey’s multiple comparison test (*p* < 0.05 vs. S100A8/A9 alone); **G**, unpaired two-tailed *t*-test (*p* < 0.05 vs. WT).

### Cigarette smoke extract aggravates atherosclerosis in *Ldlr^-/-^*mice by increasing CCR2+ monocyte-derived macrophages

Numerous studies have shown that CS exposure exacerbate atherosclerosis in mouse models^36^. Most of these studies have used whole-body or side stream smoke exposure, highlighting lung inflammation as a key contributor to the development of vascular disease^37,38^. Furthermore, exposure to CS has been shown to disrupt cholesterol metabolism, leading to elevated levels of LDL-C, total cholesterol (TC), and triglycerides (TG), along with reduced HDL-C. However, the effects of orally administered CSE on cholesterol metabolism, atherosclerosis progression, and the underlying mechanisms remain largely unclear. To explore this, we induced atherosclerosis in *Ldlr*^⁻^*^/^*^⁻^ mice by feeding them a Western diet (WD) for 14 weeks, followed by daily treatment with either vehicle or a high dose of CSE (8mg/kg) for an additional 4 weeks. As anticipated, feeding mice with a WD significantly increased body weight, blood glucose, TC and TG. However, unlike CS exposure, which is known to affect body weight and cholesterol metabolism^39^, oral administration of CSE had no impact on body weight, blood glucose, TC, or TGs (**Figure S6A through S6D**). Feeding a WD significantly increased the number of circulating monocytes and neutrophils, and these levels were further elevated in mice treated with CSE for an additional 4 weeks (**Figure 5A through 5C).** CSE administration also resulted in a significant increase in plasma S100A8/A9 levels accompanied by reduced neutrophil content in *Ldlr*^⁻^*^/^*⁻ mice **(Figure S6E)**, mirroring the effects observed in CS- or CSE-treated wild-type mice **(Figures 1J and 3G)**. These findings confirm that CS exposure, whether by inhalation or ingestion, promotes the release of S100A8/A9 from circulating neutrophils.

**Figure 5:**
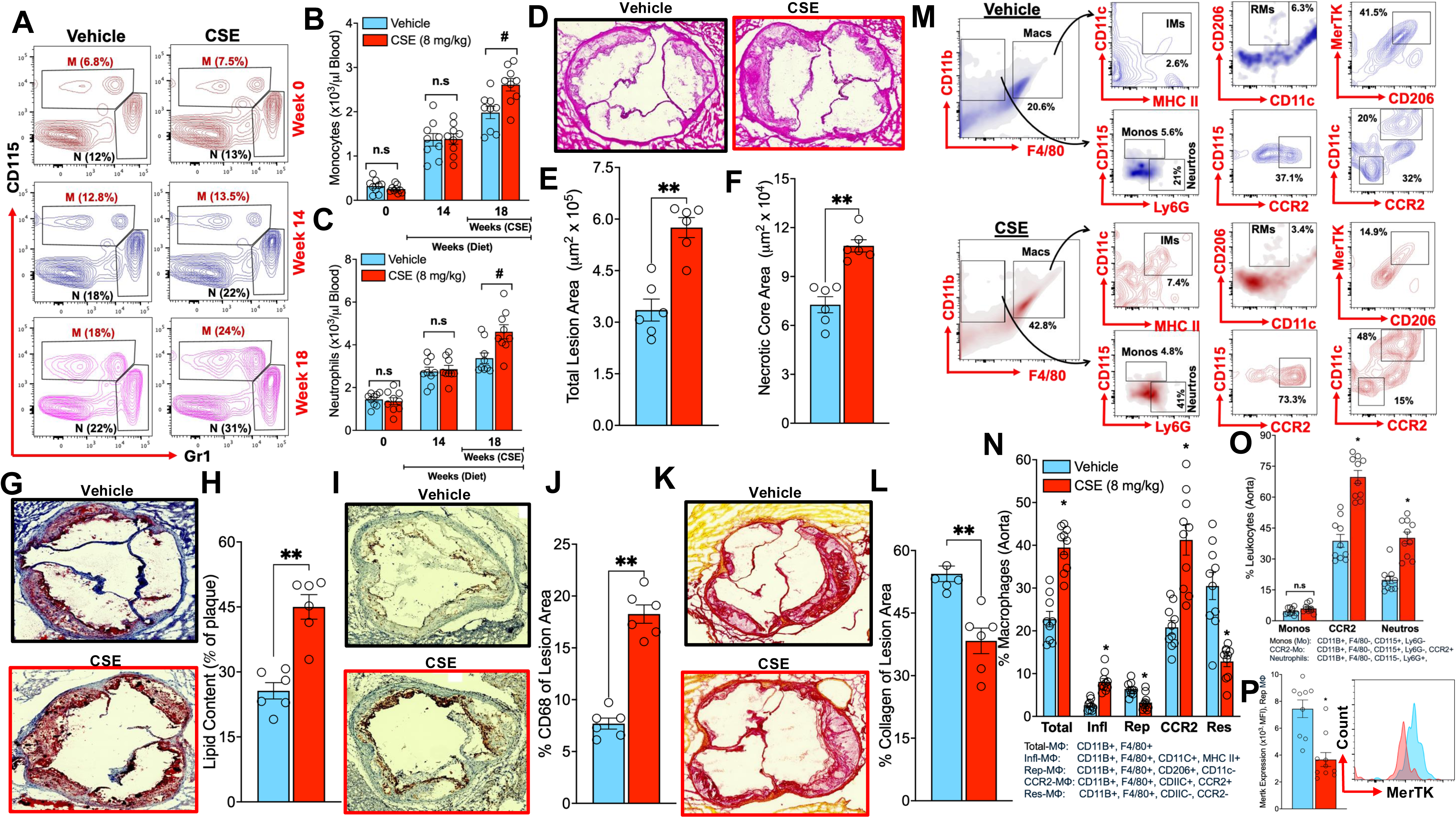
CSE aggravates atherosclerosis in *Ldlr*^⁻^*^/^*^⁻^ mice by increasing lesional CCR2⁺ macrophages and promote plaque instability. **A**, Representative flow cytometric plots depicting gating strategies for identification of monocytes and neutrophils in the blood. The values in parentheses indicate the percentage of monocytes (M) or neutrophils (N) cells in relation to CD45 cells. Quantification of circulating monocytes (**B**) and neutrophils (**C**) in *Ldlr*⁻*^/^*^⁻^ mice fed a Western diet (WD) and treated with either vehicle or CSE (8 mg/kg/day) for 4 weeks. **D**, Representative H&E-stained aortic root sections from vehicle- and CSE-treated mice, showing increased lesion size and necrotic core area in the CS**G–H**, Representative Oil Red O-stained sections (**G**) showing lipid accumulation in aortic root lesions and corresponding quantification of lipid area (**H**). CD68 macrophage immunostaining (**I**) and quantification (**J**) showing enhanced macrophage infiltration in plaques from CSE-treated mice. Picrosirius Red staining (**K**) and quantification of collagen content (**L**) in plaques, demonstrating reduced collagen deposition and features of plaque instability in the CSE group. **M**, Representative flow cytometry plots and gating strategies to identify various immune cell populations in aortic lesions. The values in parentheses indicate the percentage of either monocytes (monos), neutrophils (neutros), reparative macrophages (RMs) or inflammatory macrophages (IMs). **N**, Quantification of macrophage subtypes: inflammatory macrophages (CD11c⁺ MHC II⁺) were increased, whereas reparative macrophages (CD206⁺ CD11c⁻) were reduced in CSE-treated mice. **O**, Quantification of plaque-infiltrating neutrophils showing significant increase in the CSE group. **P**, Surface expression of MerTK on reparative macrophages was significantly downregulated following CSE treatment, indicating impaired resolution capacity. Data represent mean ± SEM. Statistical significance was determined using unpaired two-tailed *t*-tests or one-way ANOVA with Tukey’s post-hoc test as appropriate (*p* < 0.05).

Next, to investigate whether the increased number of myeloid cells in the blood was due to enhanced myelopoiesis, we examined the proliferation of HSPCs in the BM. As shown in **Figures S6F and S6G**, CSE markedly increased myelopoiesis, particularly by promoting the proliferation of CMPs and GMPs, as indicated by the rise in EdU⁺ cells. To evaluate whether the enhanced myelopoiesis induced by CSE ingestion affects atherosclerosis, we examined several markers of vascular inflammation. Analysis of H&E-stained aortic root sections revealed significantly larger atherosclerotic lesions (**Figures 5D and 5E**) and more extensive necrotic cores in CSE-treated mice (**Figures 5D and 5F**). These necrotic cores appear as pale, acellular regions (white areas) within the plaques, separated by thin bands of remaining tissue. The lesions in CSE-treated mice also displayed increased lipid content, evidenced by more extensive, red-stained areas along the inner surface and within the plaque, indicating more severe lipid accumulation compared to lesions in vehicle-treated mice (**Figure 5I and 5J**). The CSE-treated mice showed markedly higher macrophage accumulation, with approximately a 2.5-fold increase in CD68-positive staining intensity compared to vehicle-treated mice, indicating heightened vascular inflammation and immune cell infiltration. Increased macrophage burden was also associated with reduced collagen content, as evidenced by thinner and more patchy picrosirius red staining, indicating features of plaque instability (**Figure 5K and 5L)**.

While histological assessment demonstrated an increased macrophage burden, it remained unclear whether these changes were driven solely by enhanced infiltration of circulating monocytes into the plaque and/ or by impaired macrophage-mediated resolution of inflammation. To investigate this, we performed comprehensive leukocyte profiling of aortic lesions using flow cytometry. We observed a significant increase in the overall macrophage population, particularly the CD11C⁺, MHC II⁺ inflammatory macrophages, while the number of reparative macrophages (CD206⁺, CD11C⁻) was reduced in CSE-treated mice (**Figure 5M and 5N**). Although the total number of monocytes did not differ between groups, the majority of monocytes in CSE-treated lesions were CCR2⁺, suggesting increased infiltration of circulating monocytes. This shift was also accompanied by a decreased population of resident macrophages (CD11C⁻, CCR2⁻). Notably, we also observed an increased number of neutrophils within the lesions of the CSE-treated group (**Figure 5M and 5O**). Additionally, within the reparative macrophage population, surface expression of MerTK was significantly downregulated (**Figure 5P**), indicating a reduced reparative capacity of these macrophages.

### Infiltration of inflammasome-primed neutrophils aggravates atherosclerosis by impairing macrophage function

The worsening of atherosclerosis in CSE treated mice could be due to multiple factors including enhanced macrophage burden and /or impaired macrophage activity. A recent study reported that interleukin-1β (IL-1β), a product of NLRP3 inflammasome activation, suppresses MerTK and TREM2 activity in macrophages, thereby exacerbating atherosclerosis^40^. Given our observation of a significant reduction of MerTK in reparative macrophages, we hypothesized that CSE may aggravate atherosclerosis either by directly promoting IL-1β release through inflammasome priming/ activation^41,42^ or indirectly via the release of S100A8/A9 from neutrophils^35^. To investigate this, we first exposed purified monocytes and neutrophils to either CSE or S100A8/A9 and measured markers of inflammasome activation and inflammation. As shown in **Figure S7**, exposure of blood monocytes (**Figure S7A**) or neutrophils (**Figure S7B**) to CSE had no significant effect on the expression of inflammasome components or general inflammatory markers. In contrast, treatment of leukocytes with S100A8/A9 significantly increased the expression of *Nlrp3, Il1β, Tnfα,* and *Mcp1*. Similar results were observed when BM-derived macrophages (BMDMs) were exposed to either CSE or S100A8/A9 (**Figure S7C**). To determine whether the upregulation of inflammasome components in blood leukocytes leads to enhanced IL-1β release upon caspase activation, we stimulated both monocytes and neutrophils with nigericin. As expected, CSE-treated cells showed no evidence of inflammasome activation, whereas S100A8/A9-primed leukocytes exhibited a robust release of IL-1β (**Figure 6A**). These findings confirm that CSE alone does not directly induce inflammasome priming; instead, it likely depends on neutrophil-derived S100A8/A9, to promote inflammasome priming and subsequent release of IL-1β.

**Figure 6:**
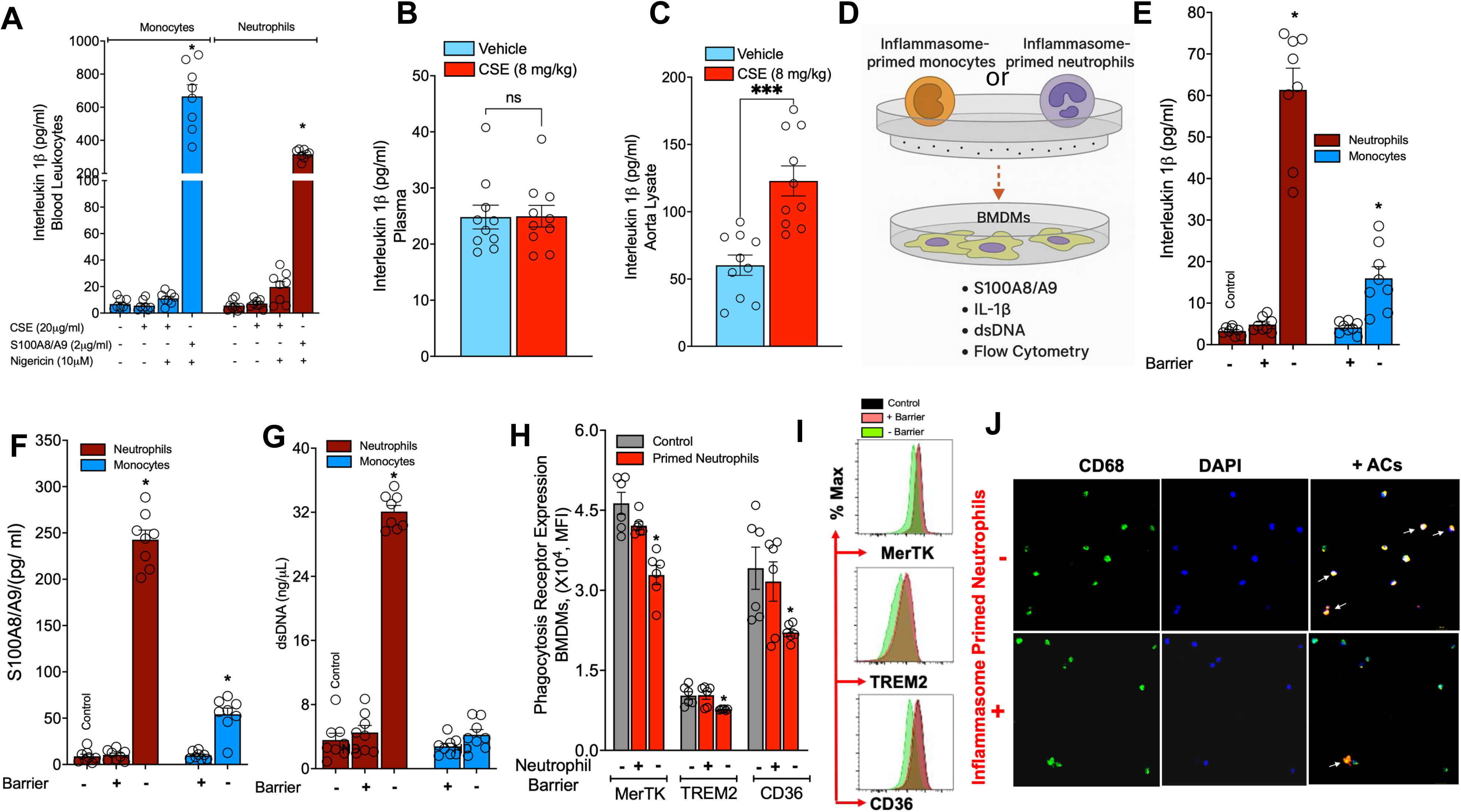
**Direct interactions between inflammasome-primed neutrophils and macrophages drive NETosis and suppress macrophage function**. **A**, Effect of CSE or S100A8/A9 on inflammasome activation (IL-1β release to nigericin) in sorted blood monocytes or neutrophils. **B**, Plasma levels of IL-1β in CSE treated *Ldlr*^⁻^*^/^*^⁻^ mice at termination. C, Protein levels of IL-1β in aortic lysates of *Ldlr*^⁻^*^/^*^⁻^ mice treated with or without CSE. **D**, Experimental setup: BMDMs were cultured in the lower chamber of a petri dish, with a 0.4 μm transwell insert placed above to permit diffusion of soluble mediators while preventing direct contact. WT monocytes or neutrophils were LPS-primed (3 h, washed) and added to the upper chamber with or without the insert. After overnight incubation, BMDM supernatants were collected for analysis of IL-1β, S100A8/A9, and dsDNA. Quantification of IL-1β (**E**), S100A8/A9 (**F**), and double stranded (ds)-DNA (**G**) released from blood neutrophils (maroon) or monocytes (blue). The presence of a transwell barrier is indicated by **+/–** symbol. The first bar in each panel represents the control condition (BMDMs exposed to media only, without added cells). **H**, Flow cytometric analysis of MerTK, TREM2, and CD36 via MFI in BMDMs exposed to LPS-primed neutrophils with or without a transwell barrier. **I**, Representative histograms showing MerTK, TREM2, and CD36 expression in control BMDMs (no added cells) and in BMDMs co-cultured with primed neutrophils in the presence or absence of a barrier. J, Representative microscopy images of BMDMs (CD68⁺) clearing CFSE (red)–labeled apoptotic cells under conditions with or without prior co-culture with inflammasome-primed neutrophils. Data represent mean ± SEM. Statistical analysis of neutrophil groups within figures **A, E, F, G and H**, one-way ANOVA with Tukey’s multiple comparison test (*p* < 0.05 vs. all other groups); For monocyte groups within figures **A, E, F, G** and **H**, unpaired two-tailed *t*-test (*p* < 0.05 vs. other groups). For **B** and **C**, unpaired two-tailed *t*-test (*p* < 0.05 vs. vehicle group, n.s, not significant)

We next measured IL-1β levels in the plasma of *Ldlr*^⁻^*^/^*^⁻^ mice and found no significant difference between PBS- and CSE-treated groups (**Figure 6B**). However, IL-1β levels were significantly elevated in the aortas of CSE-treated mice (**Figure 6C**), suggesting enhanced infiltration of inflammasome-primed monocytes and neutrophils into the lesions. These findings suggest that inflammasome-primed myeloid cells infiltrate the lesions, receive secondary activation signals from resident cells, and trigger local IL-1β release, thereby exacerbating plaque inflammation. To test this in an invitro setting, we cultured BMDMs in the bottom of a petri dish and placed a transwell insert on top, which allowed diffusion of soluble factors but prevented direct cell-cell contact. Inflammasome (i.e., LPS)-primed monocytes or neutrophils were added to the upper chamber of the insert (0.4μM) that are impermeable to them. After overnight incubation, the barriers were removed, and the supernatants from the lower chamber were collected and analyzed for IL-1β, S100A8/A9 and dsDNA levels (Study outline, **Figure 6D**). Presence of a physical barrier prevented the release of IL-1β from both monocytes and neutrophils. However, removing the barrier resulted in a dramatic release of IL-1β from neutrophils compared to monocytes (**Figure 6E**). Interestingly, the interaction between inflammasome-primed neutrophils and macrophages also resulted in a significant increase in S100A8/A9 levels (**Figure 6F**) likely due to NETosis, a form of cell death that results in large scale release of S100A8/A9 along with other cellular contents including ds-DNA (**Figure 6G**). Importantly, the BMDMs exposed to inflammasome-primed neutrophils showed a significant downregulation of various PRs including MerTK, TREM2 and CD36^43–45^ (**Figure 6H and 6I**) an effect that was also recapitulated using S100A8/A9-primed blood neutrophils (**Figure S8A**). On the other hand, the interaction of inflammasome-primed monocytes with BMDMs did not significantly impact the PRs (**Figures S8B through S8D**).

Recent studies have shown that IL-1β activates p38 MAPK to promote ADAM17-mediated cleavage of MerTK^46^, thereby linking inflammasome activation to impaired efferocytosis^40^. A similar pathway has also been implicated in TREM2 cleavage^47^, suggesting a shared mechanism through which IL-1β driven inflammatory signaling disrupts resolution programs dependent on MerTK and TREM2. To test this, we stimulated macrophages with rIL-1β and observed a significant downregulation of phagocytic receptors, including CD36, along with a marked reduction in the population of functional (MerTK⁺TREM2⁺) macrophages and acquisition of a pro-inflammatory phenotype (**Figure S8E through S8G**). Because IL-1β is elevated in the media of macrophages exposed to inflammasome-primed neutrophils (**Figure 6E**), we hypothesized that this cytokine promotes ADAM17 upregulation through p38 MAPK activation. Consistent with this, co-culture of macrophages with inflammasome-primed neutrophils led to robust induction of Adam17 and phosphorylated p38 MAPK (**Figure S8H–8I**). Similar effects were observed when macrophages were exposed to inflammasome-primed monocytes, albeit to a lesser extent, likely reflecting their moderate release of IL-1β (**Figure 6E**). Functionally, this loss of receptor expression impaired efferocytosis, as shown by reduced uptake of CFSE-labeled apoptotic cells (**Figure 6J**).

### Blocking β2 integrins averts NETosis and preserves macrophage efferocytic function

Neutrophils engage macrophages through multiple receptor-ligand interactions, most prominently via β2 integrins (Mac-1/CD11b/CD18 and LFA-1/CD11a/CD18) on neutrophils binding to ICAM-1 and related adhesion molecules on macrophages^48^. To determine whether β2 integrins drive NETosis during interactions of inflammasome-primed neutrophils with lesional macrophages, we blocked Mac-1 and LFA-1 with neutralizing antibodies and assessed macrophage receptor expression and function. In parallel, NETosis was inhibited pharmacologically to evaluate its role in IL-1β release and PR downregulation (Study Design, **Figure 7A**). Neutralization of LFA-1, either alone or combined with Mac-1, prevented neutrophil-induced downregulation of MerTK (**Figure 7B**) and TREM2 (**Figure 7C**), while CD36 expression remained preserved under all treatment conditions (**Figure 7D**). In contrast, blocking Mac-1 alone did not rescue MerTK or TREM2. Notably, inhibition of PADi4 with CI-Amidine fully preserved expression of all three PRs (**Figures 7B through 7D**). Consistent with these findings, the proportion of MerTK⁺TREM2⁺ double-positive macrophages was maintained following LFA-1 or PAD4 inhibition (**Figure 7E**), suggesting that adhesion molecules, particularly LFA-1, are essential for driving NETosis and the consequent loss of PRs. Functionally, blockade of LFA-1, combined Mac-1/LFA-1 inhibition, or PADi4 inhibition all preserved macrophage phagocytosis of apoptotic cells (**Figure 7F and 7G**). When analyzed by macrophage subpopulations by the MFI of internalized apoptotic cells, those highly expressing CD36 exhibited greater phagocytic capacity (**Figure S9A**) compared to macrophages enriched for MerTK (**Figure S9B**) or TREM2 (**Figure S9C**), although the differences among receptor-specific subsets were modest. These effects coincided with preserved expression of PRs (**Figures S9A through S9C**), reflecting effective suppression of NETosis (**Figure S9D**) and prevention of IL-1β release (**Figure S9E**) as well as S100A8/A9 (**Figure S9F**). Based on these findings, it is conceivable that inflammasome-primed neutrophils, upon entering atherosclerotic lesions during smoking, engage macrophages through adhesion receptors (e.g., LFA-1) undergo NETosis, and release IL-1β, which subsequently impairs macrophage efferocytosis by downregulating PRs.

**Figure 7:**
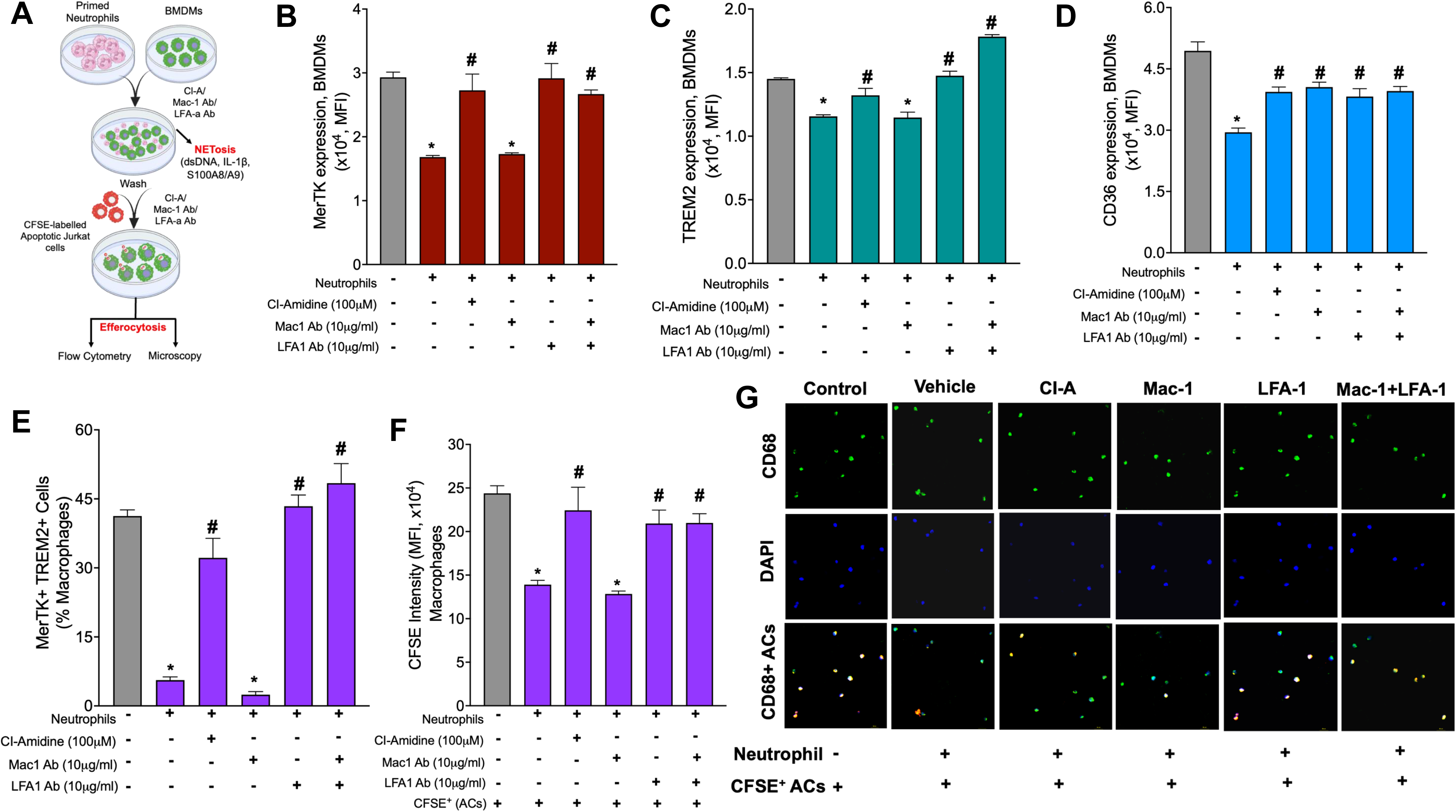
Blocking β2 integrins averts NETosis and preserves macrophage efferocytic function. **A**) Bone marrow–derived macrophages (BMDMs) were pretreated with neutralizing antibodies against Mac-1 and/or LFA-1, or with the PADi4 inhibitor CI-Amidine, followed by co-culture with inflammasome-primed neutrophils. After 3 hours, macrophages were washed thoroughly and subsequently incubated with CFSE-labeled apoptotic cells (Jurkat cells) to assess efferocytic capacity by flow cytometry and fluorescence microscopy. Flow cytometric analysis (Mean Fluorescence Intensity, MFI) of macrophage surface expression of MerTK (**B**), TREM2 (**C**), and CD36 (**D**) following co-culture with treated neutrophils. (**E**) Flow cytometric quantification of MerTK⁺TREM2⁺ double-positive macrophages. **(F**) Flow cytometric assessment of macrophage efferocytic function, measured by uptake of CFSE-labeled apoptotic cells under indicated treatment conditions. (**G**) Representative microscopy images of BMDMs (CD68⁺) clearing CFSE (red)–labeled apoptotic cells under indicated treatment conditions. Except for control conditions, BMDMs in all other groups were first co-cultured with inflammasome-primed neutrophils, then washed and incubated with CFSE-labeled apoptotic cells. Data represent mean ± SEM. Statistical analysis of figures **B, C, D, E** and **F**, one-way ANOVA with Tukey’s multiple comparison test (\**p* < 0.05 vs. control, # *p* < 0.05 vs. the neutrophil group (second bar) without inhibitors).

### Hematopoietic deletion of *S100a9* suppresses both myelopoiesis and atherosclerosis in *Ldlr*^⁻^*^/^*^⁻^ mice fed with WD

Given the central role for S100A8/A9 in mediating the pro-inflammatory effects of CS on vascular wall, we hypothesized that deletion of *S100a8/a9* in neutrophils would attenuate myelopoiesis and consequently reduce the infiltration of CCR2⁺ monocytes and inflammasome-primed neutrophils into atherosclerotic lesions. To test this, we performed BMT studies, transplanting either WT or *S100a9*^⁻^*^/^*^⁻^ BM into *Ldlr*^⁻^*^/^*^⁻^ mice, followed by treatment with CSE (Study outline, **Figure 8A**). Following successful BM reconstitution (6 weeks), the recipient mice were fed a Western diet (WD) for 14 weeks before initiating CSE treatment for an additional 4 weeks. While WD significantly increased serum glucose and lipids *S100a9* deletion had no effect on these systemic parameters (**Figure S10A through S10D**). In contrast, S100Aa9 deficiency markedly reduced the number of circulating monocytes and neutrophils (**Figure 8B and 8C**), likely by suppressed medullary myelopoiesis as evidenced by decreased proliferation of CMPs and GMPs in the BM (**Figure S10E and S10F).**

**Figure 8:**
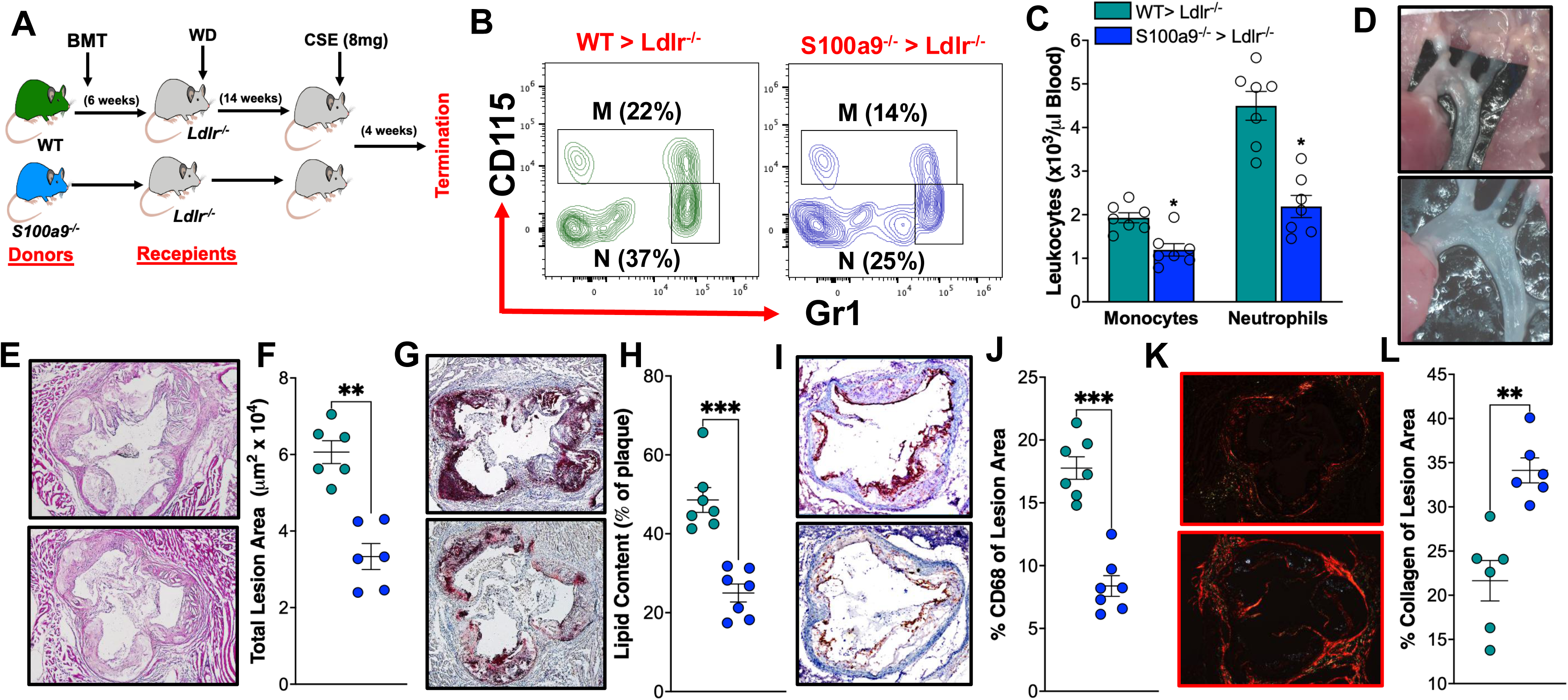
Hematopoietic deletion of *S100a9* suppresses myelopoiesis and atherosclerosis in *Ldlr*^⁻^*^/^*^⁻^ mice fed with WD. **A)**, Experimental design: WT or *S100a9*^⁻^*^/^*^⁻^ bone marrow was transplanted into *Ldlr*^⁻^*^/^*^⁻^ mice. After 6 weeks of reconstitution, recipients were fed a Western diet (WD) for 14 weeks, followed by oral gavage with CSE (8 mg/kg) for an additional 4 weeks. **B)**, Representative flow cytometric plots and gating strategy for blood monocytes and neutrophils. The numbers in parentheses indicate the percentage of monocytes (M) or neutrophils in relation to CD45. **C)** Quantification of monocytes and neutrophils at termination. **E)**, Aortic arch lesions showing severe, diffuse atherosclerosis with heavy plaque burden (top panel, *WT> Ldlr^-/-^*) while the bottom panel (*S100a9^-/-^ > Ldlr^-/-^)* shows mild condition with limited fatty streaks and preserved vessel transparency. **E)**, Representative H&E-stained cross-sections of the aortic root showing advanced atherosclerotic plaques with large necrotic cores (top) compared to more cellular, fibrous lesions with smaller necrotic areas (bottom). **F)**, Quantification of total lesion area. **G)**, Representative Oil Red O–stained sections of the aortic root showing abundant neutral lipid accumulation within large necrotic plaques (top) compared with more patchy lipid deposition throughout the intima (bottom). **H)**, Quantification of lipid content in ORO-stained images. **I)**, Representative CD68-stained cross-sections of the aortic root showing dense macrophage accumulation along the fibrous cap and shoulder regions of the plaque (top) compared with more diffuse, scattered macrophage staining throughout the lesion (bottom). **J)**, Quantification of macrophage content from CD68-stained sections. **K)**, Representative fluorescent images of aortic root sections stained for collagen (red), showing relatively thinner and less organized collagen distribution within the fibrous cap (top) compared with thicker, more extensive collagen deposition outlining the plaque and vessel wall (bottom). **L)**, Quantification of collagen content in picrosirius red stained images. Note: all top panels represent sections from WT> *Ldlr^-/-^*mice while the bottom panel shows representative images from *S100a9^-/-^* > *Ldlr^-/-^* groups. Data represent mean ± SEM. Statistical tests for **C, F, H, J** and **L**, unpaired two-tailed *t*-test (*p* < 0.05 vs. vehicle group)

Analysis of the aortic arch revealed that *S100a9*⁻*^/^*^⁻^ BMT mice developed only mild to moderate lesions, with limited white patches at the arch and branch points. In contrast, WT BMT mice exposed to CSE showed extensive, confluent lesions covering large areas of the aortic arch and branches, consistent with advanced atherosclerotic disease (**Figure 8D**). Histological analysis (H&E staining) of aortic root sections further demonstrated that WT→ *Ldlr*^⁻^*^/^*^⁻^ mice developed severe atherosclerosis, characterized by large plaques and marked lumen narrowing, whereas *S100a9*^⁻^*^/^*⁻ *Ldlr*⁻*^/^*^⁻^ mice exhibited significantly reduced plaque burden, with smaller, less occlusive lesions (**Figure 8E and 8F**). Moreover, deletion of *S100a9* substantially reduced neutral lipid accumulation (**Figure 8G and 8H**) and macrophage burden (**Figure 8I through 8K**), while preserving collagen content within the lesions (**Figure 8K and 8L**). Collectively, these findings demonstrate that loss of S100a9 in hematopoietic cells protects against CSE-induced exacerbation of atherosclerosis by limiting myelopoiesis, reducing inflammatory cell recruitment, and preserving plaque stability.

## Discussion

Our findings reveal a previously unappreciated immuno-hematopoietic pathway by which CS or its extract propels atherosclerosis independently of changes in lipid metabolism or overt lung injury. We show that CSE exposure induces oxidative stress in circulating neutrophils, prompting the release of alarmins S100A8/A9, which then engage TLR4/ RAGE receptors on BM HSPCs. This drives emergency myelopoiesis, producing elevated numbers of neutrophils and Ly6C^hi^ monocytes that infiltrate plaques. Within lesions, inflammasome-primed neutrophils undergo NETosis and liberate IL-1β, causing downregulation of key PRs including MerTK, TREM2 and CD36 on macrophages, compromising their efferocytosis function and exacerbating plaque progression. Hematopoietic deletion of *S100a9* disrupts this axis, markedly reducing myelopoiesis, lesion inflammation, and plaque vulnerability even under continued CSE exposure. These findings extend the current understanding of smoking-induced cardiovascular disease.

Elevated S100A8/A9 levels in humans have long been associated with neutrophil counts, smoking status, and CVD risk^49,50^. Multiple studies demonstrate that CS exposure markedly increases S100A8/A9 expression across neutrophil subsets in both BM and blood, reprogramming granulopoiesis to generate neutrophils transcriptionally primed with high S100A8/A9^33^. Importantly, S100A8/A9 function not only as biomarkers but also as active mediators that promote leukocyte recruitment and myeloid expansion^27,35,51,52^. These findings suggest that a systemic expansion of alarmin-rich, inflammasome-competent neutrophils are capable of amplifying vascular inflammation. Our work builds on this foundation by showing that CS acts as a potent inducer of neutrophil oxidative stress and S100A8/A9 release, thereby translating a common environmental insult into systemic hematopoietic activation. While the concept of heightened myelopoiesis accelerating atherosclerosis is increasingly recognized ^53,54^, the upstream stimuli in smokers have remained unclear. By identifying neutrophil-derived S100A8/A9 as a central signaling hub linking toxic environmental exposure with BM activation, our study fills a critical mechanistic gap. Moreover, whereas previous models largely attributed CS’s atherogenic effects to dyslipidemia or endothelial dysfunction^55,56^, our data reveal a distinctly inflammation-driven pathway: CS exposure promotes systemic inflammation and accelerates plaque progression without perturbing lipid metabolism, representing a significant shift from prevailing paradigms.

Our BMT studies provide compelling functional evidence that S100A8/A9 plays a pivotal role in atherosclerotic plaque progression. Specifically, the loss of *S100a9* in the hematopoietic compartment led to reductions in circulating myeloid cells, diminished lipid accumulation and macrophage burden within the lesions, and preservation of fibrous collagen, the features that collectively suggest enhanced plaque stability. These findings elevate S100A8/A9 from a mere biomarker of inflammation to a *bona fide* therapeutic target with the potential to intercept upstream drivers of vascular inflammation. Targeting this axis through neutralizing antibodies, receptor blockade (e.g., TLR4 or RAGE), or small-molecule inhibitors could disrupt the deleterious feed-forward loop between neutrophil-derived alarmins and myeloid-driven inflammation. Indeed, preclinical studies have demonstrated that inhibition of S100A8/A9 can attenuate atherosclerosis progression ^57^, as well as reduce post-myocardial infarction (MI) remodeling and adverse outcomes post-MI remodeling^35,58,59^. These findings underscore the broader therapeutic relevance of this axis beyond atherosclerosis alone.

The role of inflammasome-primed neutrophils in amplifying lesion inflammation reveals a previously underrecognized axis of innate immune crosstalk in cigarette smokers. These neutrophils, through NETosis and localized IL-1β release, can profoundly impair the reparative functions of macrophages. Evidence from diabetic mouse models shows that NETs exacerbate macrophage-driven inflammation and hinder atherosclerosis resolution, creating a sustained pro-inflammatory environment within plaques^60^ . Furthermore, IL-1β generated downstream of inflammasome activation not only perpetuates inflammatory signaling but also directly triggers cleavage of the efferocytosis receptor MerTK and TREM2 via a p38-ADAM17-mediated pathway, thereby undermining macrophage efferocytosis and plaque resolution^40^. The resulting attenuation of surface receptors such as MerTK and TREM2 is likely a critical mechanism by which macrophages fail to clear apoptotic cells efficiently and fostering vulnerable plaque morphology.

However, it’s important to acknowledge key limitations. *First*, while our findings implicate predominantly neutrophil-driven IL-1β in macrophage dysfunction, the specific macrophage subsets affected remain undefined. Atherosclerotic lesions contain resident TREM2^hi^ reparative macrophages, CCR2⁺ monocyte-derived inflammatory macrophages, and foamy lipid-processing variants each contributing differently to lesion dynamics^61^. Our data do not resolve whether reparative macrophages versus inflammatory CCR2⁺ counterparts are most susceptible to IL-1β-mediated suppression. Subset-specific lineage tracing, conditional knockouts, and single-cell RNA sequencing are needed to clarify which macrophage compartments are most disturbed by inflammasome activation. *Second*, although IL-1β is a key driver, other inflammatory mediators may also contribute especially given that NETosis is the major source of IL-1β in the lesions. NET-derived histones and proteases can exacerbate plaque inflammation independently of IL-1β^62^, and S100A8/A9 may further drive NET formation via RAGE signaling^63^. Future work targeting NETosis or RAGE should disentangle these parallel and potentially synergistic pathways. *Third*, the complexity of CS itself is a significant confounder. CS comprises thousands of constituents including nicotine, acrolein, aldehydes, and fine particulate matter (PM₂.₅), many of which can individually provoke oxidative stress, inflammatory activation, or hematopoietic responses. For example, nicotine and PM_2.5_ exposure activates the NLRP3 inflammasome^64,65^. Acrolein promotes oxidative stress and destabilize atherosclerotic plaques^66^. Without fractionation studies or model exposures to isolated constituents, we cannot pinpoint which components drive neutrophil priming and S100A8/A9 release. Future experimental models using purified components and real-world smoke exposures are needed to resolve this question. *Fourth*, although we show that NETosis of neutrophils downregulates macrophage PRs, the mechanism underlying CD36 loss remains unclear. Unlike MerTK and TREM2, where IL-1β–p38–ADAM17 pathways are implicated, it is unknown whether CD36 downregulation is mediated directly by IL-1β or through other cytokines, signaling cascades, or proteolytic mechanisms. Dissecting this will require targeted blocking studies and transcriptomic profiling to determine whether CD36 is suppressed transcriptionally, shed proteolytically, or regulated through alternative inflammatory mediators. Despite these limitations, our findings hold promising translational implications. S100A8/A9 emerges as both a biomarker and a therapeutic target. Plasma levels of S100A8/A9 predict adverse outcomes in smokers and patients with coronary artery disease. Strategically, targeting both the upstream S100A8/A9 axis and downstream IL-1β effects via NLRP3 or IL-1β inhibitors may offer synergistic control of smoking-induced vascular inflammation.

In summary, our studies reveal a neutrophil-BM-vessel wall signaling axis, mediated by S100A8/A9, as a fundamental pathway by which CS accelerates atherosclerosis independently of lipid profiles. This axis encompasses neutrophil oxidative activation, emergency myelopoiesis, lesion infiltration, macrophage impairment, and plaque vulnerability. Disrupting this pathway offers new opportunities to reduce cardiovascular risk among smokers and users of alternative tobacco products, potentially shifting the therapeutic paradigm to include immunometabolic-targeted strategies. Furthermore, evolution in nicotine delivery methods including e-cigarettes and oral nicotine products adds urgency. Our demonstration that oral CSE replicates smoking-induced inflammation suggests that systemic neutrophil activation and myelopoietic disruption may occur even without lung injury, calling for caution in the assumption that non-inhalational forms of nicotine are innocuous.

## Author’s Contribution

D.C., N.N., R.J., and P.N., conceived and designed the study, analyzed the data, created figures, and wrote the main manuscript that was revised and approved by all authors. B.A., K.P., A.D., R.S., M.L., performed animal studies, cell culture experiments, flow cytometry, ELISAs, WBs, RT-PCRs, and histological analysis. A.M contributed to manuscript writing, editing, discussion and critical inputs.

## Supplemental Materials

The online-only data supplement contains Expanded materials and methods, Supplemental figures S1-S10 and Supplemental figure legends.

## Non-Standard Abbreviations

AM: Alveolar macrophages
BALF: bronchoalveolar lavage fluid
BMDMs: Bone marrow-derived macrophages
BM: Bone marrow
BMT: Bone marrow transplant
CCR2: C-C chemokine receptor type 2
CS: Cigarette smoking/ smoke
CSE: Cigarette smoke extract
CMP: Common myeloid progenitor
CVD: Cardiovascular disease
DC: Dendritic cells
DPI: Diphenyleneiodonium
dsDNA: Double-stranded DNA
ECs: Endothelial cells
GMP: Granulocyte-monocyte progenitor
HDL: High-density lipoprotein
HSPCs: Hematopoietic stem and progenitor cells
IL-1β: Interleukin 1 beta
IM: Interstitial macrophages
LDL: Low-density lipoprotein
Ldlr^-/-^: Low-density lipoprotein receptor-deficient
MHC II^+^: Major Histocompatibility Complex Class II positive
MFI: Mean fluorescence intensity
MerTK: Mer tyrosine kinase
NETosis: Neutrophil extracellular trap formation
NETs: Neutrophil extracellular traps
PMA: Phorbol myristate acetate
RAGE: Receptor for advanced glycation endproducts
S100A8/A9: S100 calcium-binding protein A8/A9
TG: Triglycerides
TC: Total cholesterol
TLR4: Toll like receptor -4
TREM2: Triggering receptor expressed on myeloid cells 2
VLDL: Very-low-density lipoprotein
WD: Western diet
WT: Wild type

## References

1. Makrygiannakis D, Hermansson M, Ulfgren AK, Nicholas AP, Zendman AJ, Eklund A, Grunewald J, Skold CM, Klareskog L, Catrina AI. Smoking increases peptidylarginine deiminase 2 enzyme expression in human lungs and increases citrullination in BAL cells. Ann Rheum Dis. 2008;67:1488–1492. doi: 10.1136/ard.2007.075192

2. Sopori M. Effects of cigarette smoke on the immune system. Nat Rev Immunol. 2002;2:372–377. doi: 10.1038/nri803

3. Messner B, Bernhard D. Smoking and cardiovascular disease: mechanisms of endothelial dysfunction and early atherogenesis. Arterioscler Thromb Vasc Biol. 2014;34:509–515. doi: 10.1161/ATVBAHA.113.300156

4. Katanoda K, Yako-Suketomo H. Mortality attributable to tobacco by selected countries based on the WHO Global Report. Jpn J Clin Oncol. 2012;42:561–562. doi: 10.1093/jjco/hys083

5. Dahdah A, Jaggers RM, Sreejit G, Johnson J, Kanuri B, Murphy AJ, Nagareddy PR. Immunological Insights into Cigarette Smoking-Induced Cardiovascular Disease Risk. Cells. 2022;11. doi: 10.3390/cells11203190

6. Lavi S, Prasad A, Yang EH, Mathew V, Simari RD, Rihal CS, Lerman LO, Lerman A. Smoking is associated with epicardial coronary endothelial dysfunction and elevated white blood cell count in patients with chest pain and early coronary artery disease. Circulation. 2007;115:2621–2627. doi: 10.1161/CIRCULATIONAHA.106.641654

7. Basilico P, Cremona TP, Oevermann A, Piersigilli A, Benarafa C. Increased Myeloid Cell Production and Lung Bacterial Clearance in Mice Exposed to Cigarette Smoke. Am J Respir Cell Mol Biol. 2016;54:424–435. doi: 10.1165/rcmb.2015-0017OC

8. Celermajer DS, Sorensen KE, Gooch VM, Spiegelhalter DJ, Miller OI, Sullivan ID, Lloyd JK, Deanfield JE. Non-invasive detection of endothelial dysfunction in children and adults at risk of atherosclerosis. Lancet. 1992;340:1111–1115. doi: 10.1016/0140-6736(92)93147-f

9. Centner AM, Cullen AE, Khalili L, Ukhanov V, Hill S, Deitado R, Hwang HS, Azeez T, La Favor JD, Laitano O, et al. The Role of Sex in the Effects of Smoking and Nicotine on Cardiovascular Function, Atherosclerosis, and Inflammation. Nicotine Tob Res. 2025;27:1116–1126. doi: 10.1093/ntr/ntae274

10. Craig WY, Palomaki GE, Haddow JE. Cigarette smoking and serum lipid and lipoprotein concentrations: an analysis of published data. BMJ. 1989;298:784–788. doi: 10.1136/bmj.298.6676.784

11. Neufeld EJ, Mietus-Snyder M, Beiser AS, Baker AL, Newburger JW. Passive cigarette smoking and reduced HDL cholesterol levels in children with high-risk lipid profiles. Circulation. 1997;96:1403–1407. doi: 10.1161/01.cir.96.5.1403

12. Freedman DS, Srinivasan SR, Shear CL, Hunter SM, Croft JB, Webber LS, Berenson GS. Cigarette smoking initiation and longitudinal changes in serum lipids and lipoproteins in early adulthood: the Bogalusa Heart Study. Am J Epidemiol. 1986;124:207–219. doi: 10.1093/oxfordjournals.aje.a114379

13. Morrow JD, Frei B, Longmire AW, Gaziano JM, Lynch SM, Shyr Y, Strauss WE, Oates JA, Roberts LJ, 2nd. Increase in circulating products of lipid peroxidation (F2-isoprostanes) in smokers. Smoking as a cause of oxidative damage. N Engl J Med. 1995;332:1198–1203. doi: 10.1056/NEJM199505043321804

14. Cavusoglu Y, Timuralp B, Us T, Akgun Y, Kudaiberdieva G, Gorenek B, Unalir A, Goktekin O, Ata N. Cigarette smoking increases plasma concentrations of vascular cell adhesion molecule-1 in patients with coronary artery disease. Angiology. 2004;55:397–402. doi: 10.1177/000331970405500406

15. Pilz H, Oguogho A, Chehne F, Lupattelli G, Palumbo B, Sinzinger H. Quitting cigarette smoking results in a fast improvement of in vivo oxidation injury (determined via plasma, serum and urinary isoprostane). Thromb Res. 2000;99:209–221. doi: 10.1016/s0049-3848(00)00249-8

16. Reilly M, Delanty N, Lawson JA, FitzGerald GA. Modulation of oxidant stress in vivo in chronic cigarette smokers. Circulation. 1996;94:19–25. doi: 10.1161/01.cir.94.1.19

17. Solak ZA, Kabaroglu C, Cok G, Parildar Z, Bayindir U, Ozmen D, Bayindir O. Effect of different levels of cigarette smoking on lipid peroxidation, glutathione enzymes and paraoxonase 1 activity in healthy people. Clin Exp Med. 2005;5:99–105. doi: 10.1007/s10238-005-0072-5

18. Barcelo B, Pons J, Ferrer JM, Sauleda J, Fuster A, Agusti AG. Phenotypic characterisation of T-lymphocytes in COPD: abnormal CD4+CD25+ regulatory T-lymphocyte response to tobacco smoking. Eur Respir J. 2008;31:555–562. doi: 10.1183/09031936.00010407

19. Chiappori A, Folli C, Balbi F, Caci E, Riccio AM, De Ferrari L, Melioli G, Braido F, Canonica GW. CD4(+)CD25(high)CD127(-) regulatory T-cells in COPD: smoke and drugs effect. World Allergy Organ J. 2016;9:5. doi: 10.1186/s40413-016-0095-2

20. Brandsma CA, Hylkema MN, Geerlings M, van Geffen WH, Postma DS, Timens W, Kerstjens HA. Increased levels of (class switched) memory B cells in peripheral blood of current smokers. Respir Res. 2009;10:108. doi: 10.1186/1465-9921-10-108

21. Brandsma CA, Kerstjens HA, van Geffen WH, Geerlings M, Postma DS, Hylkema MN, Timens W. Differential switching to IgG and IgA in active smoking COPD patients and healthy controls. Eur Respir J. 2012;40:313–321. doi: 10.1183/09031936.00011211

22. Arnson Y, Shoenfeld Y, Amital H. Effects of tobacco smoke on immunity, inflammation and autoimmunity. J Autoimmun. 2010;34:J258–265. doi: 10.1016/j.jaut.2009.12.003

23. Hoenderdos K, Condliffe A. The neutrophil in chronic obstructive pulmonary disease. Am J Respir Cell Mol Biol. 2013;48:531–539. doi: 10.1165/rcmb.2012-0492TR

24. Mills KH, Dunne A. Immune modulation: IL-1, master mediator or initiator of inflammation. Nat Med. 2009;15:1363–1364. doi: 10.1038/nm1209-1363

25. Striz I, Brabcova E, Kolesar L, Sekerkova A. Cytokine networking of innate immunity cells: a potential target of therapy. Clin Sci (Lond). 2014;126:593–612. doi: 10.1042/CS20130497

26. Eltom S, Belvisi MG, Stevenson CS, Maher SA, Dubuis E, Fitzgerald KA, Birrell MA. Role of the inflammasome-caspase1/11-IL-1/18 axis in cigarette smoke driven airway inflammation: an insight into the pathogenesis of COPD. PLoS One. 2014;9:e112829. doi: 10.1371/journal.pone.0112829

27. Nagareddy PR, Murphy AJ, Stirzaker RA, Hu Y, Yu S, Miller RG, Ramkhelawon B, Distel E, Westerterp M, Huang LS, et al. Hyperglycemia promotes myelopoiesis and impairs the resolution of atherosclerosis. Cell Metab. 2013;17:695–708. doi: 10.1016/j.cmet.2013.04.001

28. Combadiere C, Potteaux S, Rodero M, Simon T, Pezard A, Esposito B, Merval R, Proudfoot A, Tedgui A, Mallat Z. Combined inhibition of CCL2, CX3CR1, and CCR5 abrogates Ly6C(hi) and Ly6C(lo) monocytosis and almost abolishes atherosclerosis in hypercholesterolemic mice. Circulation. 2008;117:1649–1657. doi: CIRCULATIONAHA.107.745091 [pii] 10.1161/CIRCULATIONAHA.107.745091

29. Murphy AJ, Akhtari M, Tolani S, Pagler T, Bijl N, Kuo CL, Wang M, Sanson M, Abramowicz S, Welch C, et al. ApoE regulates hematopoietic stem cell proliferation, monocytosis, and monocyte accumulation in atherosclerotic lesions in mice. The Journal of clinical investigation. 2011;121:4138–4149. doi: 10.1172/JCI57559

30. Rajavashisth T, Qiao JH, Tripathi S, Tripathi J, Mishra N, Hua M, Wang XP, Loussararian A, Clinton S, Libby P, et al. Heterozygous osteopetrotic (op) mutation reduces atherosclerosis in LDL receptor-deficient mice. The Journal of clinical investigation. 1998;101:2702–2710. doi: 10.1172/JCI119891

31. Swirski FK, Libby P, Aikawa E, Alcaide P, Luscinskas FW, Weissleder R, Pittet MJ. Ly-6Chi monocytes dominate hypercholesterolemia-associated monocytosis and give rise to macrophages in atheromata. The Journal of clinical investigation. 2007;117:195–205. doi: 10.1172/JCI29950

32. Sreejit G, Abdel-Latif A, Athmanathan B, Annabathula R, Dhyani A, Noothi SK, Quaife-Ryan GA, Al-Sharea A, Pernes G, Dragoljevic D, et al. Neutrophil-Derived S100A8/A9 Amplify Granulopoiesis Following Myocardial Infarction. Circulation. 2020;141:1080–1094. doi: 10.1161/CIRCULATIONAHA.119.043833

33. Kapellos TS, Bassler K, Fujii W, Nalkurthi C, Schaar AC, Bonaguro L, Pecht T, Galvao I, Agrawal S, Saglam A, et al. Systemic alterations in neutrophils and their precursors in early-stage chronic obstructive pulmonary disease. Cell Rep. 2023;42:112525. doi: 10.1016/j.celrep.2023.112525

34. Tardif MR, Chapeton-Montes JA, Posvandzic A, Page N, Gilbert C, Tessier PA. Secretion of S100A8, S100A9, and S100A12 by Neutrophils Involves Reactive Oxygen Species and Potassium Efflux. J Immunol Res. 2015;2015:296149. doi: 10.1155/2015/296149

35. Sreejit G, Abdel-Latif A, Athmanathan B, Annabathula R, Dhyani A, Noothi SK, Quaife-Ryan GA, Al-Sharea A, Pernes G, Dragoljevic D, et al. Neutrophil-Derived S100A8/A9 Amplify Granulopoiesis After Myocardial Infarction. Circulation. 2020;141:1080–1094. doi: 10.1161/CIRCULATIONAHA.119.043833

36. Thayaparan D, Emoto T, Khan AB, Besla R, Hamidzada H, El-Maklizi M, Sivasubramaniyam T, Vohra S, Hagerman A, Nejat S, et al. Endothelial dysfunction drives atherosclerotic plaque macrophage-dependent abdominal aortic aneurysm formation. Nat Immunol. 2025;26:706–721. doi: 10.1038/s41590-025-02132-8

37. Lau PP, Li L, Merched AJ, Zhang AL, Ko KW, Chan L. Nicotine induces proinflammatory responses in macrophages and the aorta leading to acceleration of atherosclerosis in low-density lipoprotein receptor(-/-) mice. Arterioscler Thromb Vasc Biol. 2006;26:143–149. doi: 10.1161/01.ATV.0000193510.19000.10

38. Mullick AE, Tobias PS, Curtiss LK. Modulation of atherosclerosis in mice by Toll-like receptor 2. The Journal of clinical investigation. 2005;115:3149–3156. doi: 10.1172/JCI25482

39. Chen H, Hansen MJ, Jones JE, Vlahos R, Anderson GP, Morris MJ. Detrimental metabolic effects of combining long-term cigarette smoke exposure and high-fat diet in mice. Am J Physiol Endocrinol Metab. 2007;293:E1564–1571. doi: 10.1152/ajpendo.00442.2007

40. Liu W, Hardaway BD, Kim E, Pauli J, Wettich JL, Yalcinkaya M, Hsu CC, Xiao T, Reilly MP, Tabas I, et al. Inflammatory crosstalk impairs phagocytic receptors and aggravates atherosclerosis in clonal hematopoiesis in mice. The Journal of clinical investigation. 2024;135. doi: 10.1172/JCI182939

41. Dino P, Giuffre MR, Buscetta M, Di Vincenzo S, La Mensa A, Cristaldi M, Bucchieri F, Lo Iacono G, Bertani A, Pace E, et al. Release of IL-1beta and IL-18 in human primary bronchial epithelial cells exposed to cigarette smoke is independent of NLRP3. Eur J Immunol. 2024;54:e2451053. doi: 10.1002/eji.202451053

42. Mehta S, Srivastava N, Bhatia A, Dhawan V. Exposure of cigarette smoke condensate activates NLRP3 inflammasome in vitro and in vivo: A connotation of innate immunity and atherosclerosis. Int Immunopharmacol. 2020;84:106561. doi: 10.1016/j.intimp.2020.106561

43. Jaitin DA, Adlung L, Thaiss CA, Weiner A, Li B, Descamps H, Lundgren P, Bleriot C, Liu Z, Deczkowska A, et al. Lipid-Associated Macrophages Control Metabolic Homeostasis in a Trem2-Dependent Manner. Cell. 2019;178:686–698 e614. doi: 10.1016/j.cell.2019.05.054

44. Patterson MT, Firulyova MM, Xu Y, Hillman H, Bishop C, Zhu A, Hickok GH, Schrank PR, Ronayne CE, Caillot Z, et al. Trem2 promotes foamy macrophage lipid uptake and survival in atherosclerosis. Nat Cardiovasc Res. 2023;2:1015–1031. doi: 10.1038/s44161-023-00354-3

45. Tian K, Xu Y, Sahebkar A, Xu S. CD36 in Atherosclerosis: Pathophysiological Mechanisms and Therapeutic Implications. Curr Atheroscler Rep. 2020;22:59. doi: 10.1007/s11883-020-00870-8

46. Thorp E, Vaisar T, Subramanian M, Mautner L, Blobel C, Tabas I. Shedding of the Mer tyrosine kinase receptor is mediated by ADAM17 protein through a pathway involving reactive oxygen species, protein kinase Cdelta, and p38 mitogen-activated protein kinase (MAPK). J Biol Chem. 2011;286:33335–33344. doi: 10.1074/jbc.M111.263020

47. Wang X, He Q, Zhou C, Xu Y, Liu D, Fujiwara N, Kubota N, Click A, Henderson P, Vancil J, et al. Prolonged hypernutrition impairs TREM2-dependent efferocytosis to license chronic liver inflammation and NASH development. Immunity. 2023;56:58–77 e11. doi: 10.1016/j.immuni.2022.11.013

48. Zhang X, Kang Z, Yin D, Gao J. Role of neutrophils in different stages of atherosclerosis. Innate Immun. 2023;29:97–109. doi: 10.1177/17534259231189195

49. Cotoi OS, Duner P, Ko N, Hedblad B, Nilsson J, Bjorkbacka H, Schiopu A. Plasma S100A8/A9 correlates with blood neutrophil counts, traditional risk factors, and cardiovascular disease in middle-aged healthy individuals. Arterioscler Thromb Vasc Biol. 2014;34:202–210. doi: 10.1161/ATVBAHA.113.302432

50. Schiopu A, Cotoi OS. S100A8 and S100A9: DAMPs at the crossroads between innate immunity, traditional risk factors, and cardiovascular disease. Mediators Inflamm. 2013;2013:828354. doi: 10.1155/2013/828354

51. Nagareddy PR, Kraakman M, Masters SL, Stirzaker RA, Gorman DJ, Grant RW, Dragoljevic D, Hong ES, Abdel-Latif A, Smyth SS, et al. Adipose tissue macrophages promote myelopoiesis and monocytosis in obesity. Cell Metab. 2014;19:821–835. doi: 10.1016/j.cmet.2014.03.029

52. Wang S, Song R, Wang Z, Jing Z, Wang S, Ma J. S100A8/A9 in Inflammation. Front Immunol. 2018;9:1298. doi: 10.3389/fimmu.2018.01298

53. Dragoljevic D, Westerterp M, Veiga CB, Nagareddy P, Murphy AJ. Disordered haematopoiesis and cardiovascular disease: a focus on myelopoiesis. Clin Sci (Lond). 2018;132:1889–1899. doi: 10.1042/CS20180111

54. Flynn MC, Pernes G, Lee MKS, Nagareddy PR, Murphy AJ. Monocytes, Macrophages, and Metabolic Disease in Atherosclerosis. Front Pharmacol. 2019;10:666. doi: 10.3389/fphar.2019.00666

55. Ishida M, Sakai C, Kobayashi Y, Ishida T. Cigarette Smoking and Atherosclerotic Cardiovascular Disease. J Atheroscler Thromb. 2024;31:189–200. doi: 10.5551/jat.RV22015

56. Csordas A, Bernhard D. The biology behind the atherothrombotic effects of cigarette smoke. Nat Rev Cardiol. 2013;10:219–230. doi: 10.1038/nrcardio.2013.8

57. Kraakman MJ, Lee MK, Al-Sharea A, Dragoljevic D, Barrett TJ, Montenont E, Basu D, Heywood S, Kammoun HL, Flynn M, et al. Neutrophil-derived S100 calcium-binding proteins A8/A9 promote reticulated thrombocytosis and atherogenesis in diabetes. The Journal of clinical investigation. 2017;127:2133–2147. doi: 10.1172/JCI92450

58. Jakobsson G, Papareddy P, Andersson H, Mulholland M, Bhongir R, Ljungcrantz I, Engelbertsen D, Bjorkbacka H, Nilsson J, Manea A, et al. Therapeutic S100A8/A9 blockade inhibits myocardial and systemic inflammation and mitigates sepsis-induced myocardial dysfunction. Crit Care. 2023;27:374. doi: 10.1186/s13054-023-04652-x

59. Marinkovic G, Grauen Larsen H, Yndigegn T, Szabo IA, Mares RG, de Camp L, Weiland M, Tomas L, Goncalves I, Nilsson J, et al. Inhibition of pro-inflammatory myeloid cell responses by short-term S100A9 blockade improves cardiac function after myocardial infarction. Eur Heart J. 2019;40:2713–2723. doi: 10.1093/eurheartj/ehz461

60. Josefs T, Barrett TJ, Brown EJ, Quezada A, Wu X, Voisin M, Amengual J, Fisher EA. Neutrophil extracellular traps promote macrophage inflammation and impair atherosclerosis resolution in diabetic mice. JCI Insight. 2020;5. doi: 10.1172/jci.insight.134796

61. Willemsen L, de Winther MP. Macrophage subsets in atherosclerosis as defined by single-cell technologies. J Pathol. 2020;250:705–714. doi: 10.1002/path.5392

62. Silvestre-Roig C, Braster Q, Wichapong K, Lee EY, Teulon JM, Berrebeh N, Winter J, Adrover JM, Santos GS, Froese A, et al. Externalized histone H4 orchestrates chronic inflammation by inducing lytic cell death. Nature. 2019;569:236–240. doi: 10.1038/s41586-019-1167-6

63. Du F, Ding Z, Ronnow CF, Rahman M, Schiopu A, Thorlacius H. S100A9 induces reactive oxygen species-dependent formation of neutrophil extracellular traps in abdominal sepsis. Exp Cell Res. 2022;421:113405. doi: 10.1016/j.yexcr.2022.113405

64. Hu T, Zhu P, Liu Y, Zhu H, Geng J, Wang B, Yuan G, Peng Y, Xu B. PM2.5 induces endothelial dysfunction via activating NLRP3 inflammasome. Environ Toxicol. 2021;36:1886–1893. doi: 10.1002/tox.23309

65. Wu X, Zhang H, Qi W, Zhang Y, Li J, Li Z, Lin Y, Bai X, Liu X, Chen X, et al. Nicotine promotes atherosclerosis via ROS-NLRP3-mediated endothelial cell pyroptosis. Cell Death Dis. 2018;9:171. doi: 10.1038/s41419-017-0257-3

66. O’Toole TE, Zheng YT, Hellmann J, Conklin DJ, Barski O, Bhatnagar A. Acrolein activates matrix metalloproteinases by increasing reactive oxygen species in macrophages. Toxicol Appl Pharmacol. 2009;236:194–201. doi: 10.1016/j.taap.2009.01.024

