## Supplementary figures and images for "Cigarette smoke aggravates atherosclerosis by promoting the infiltration of inflammasome-primed neutrophils and disrupting macrophage function in lesions"

### Supplemental Figure-1

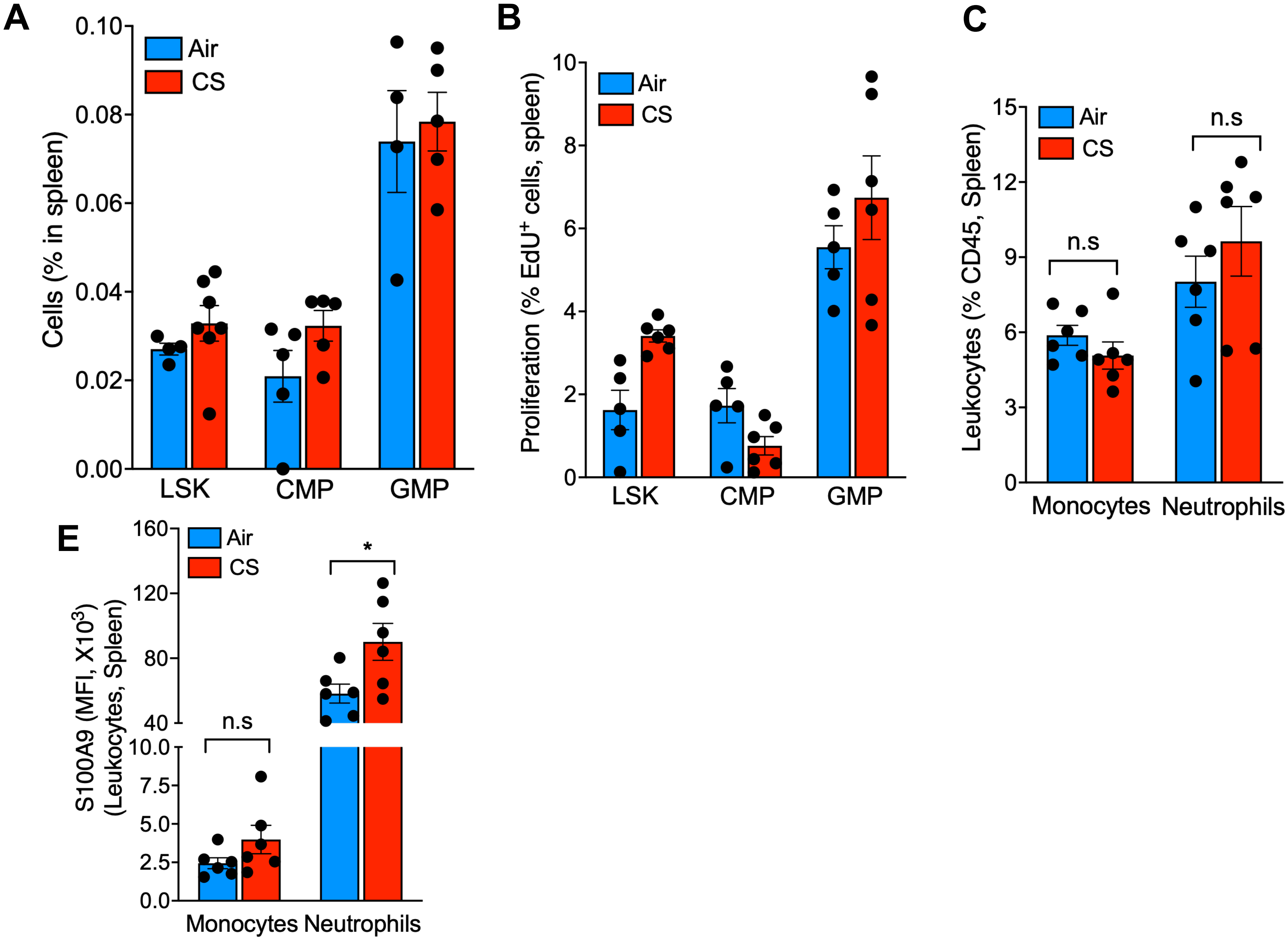

### Supplemental Figure-2

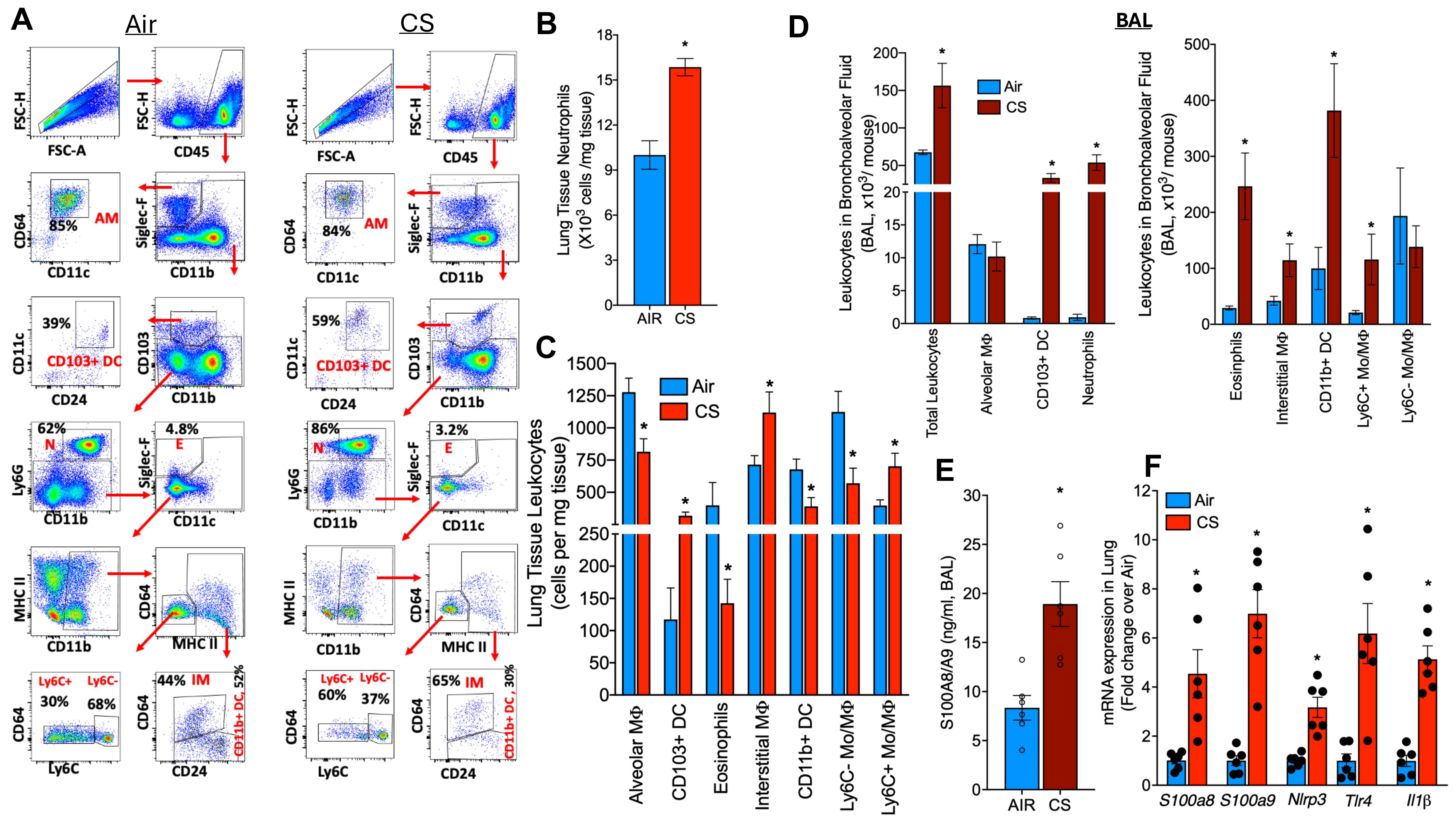

### Supplemental Figure-3

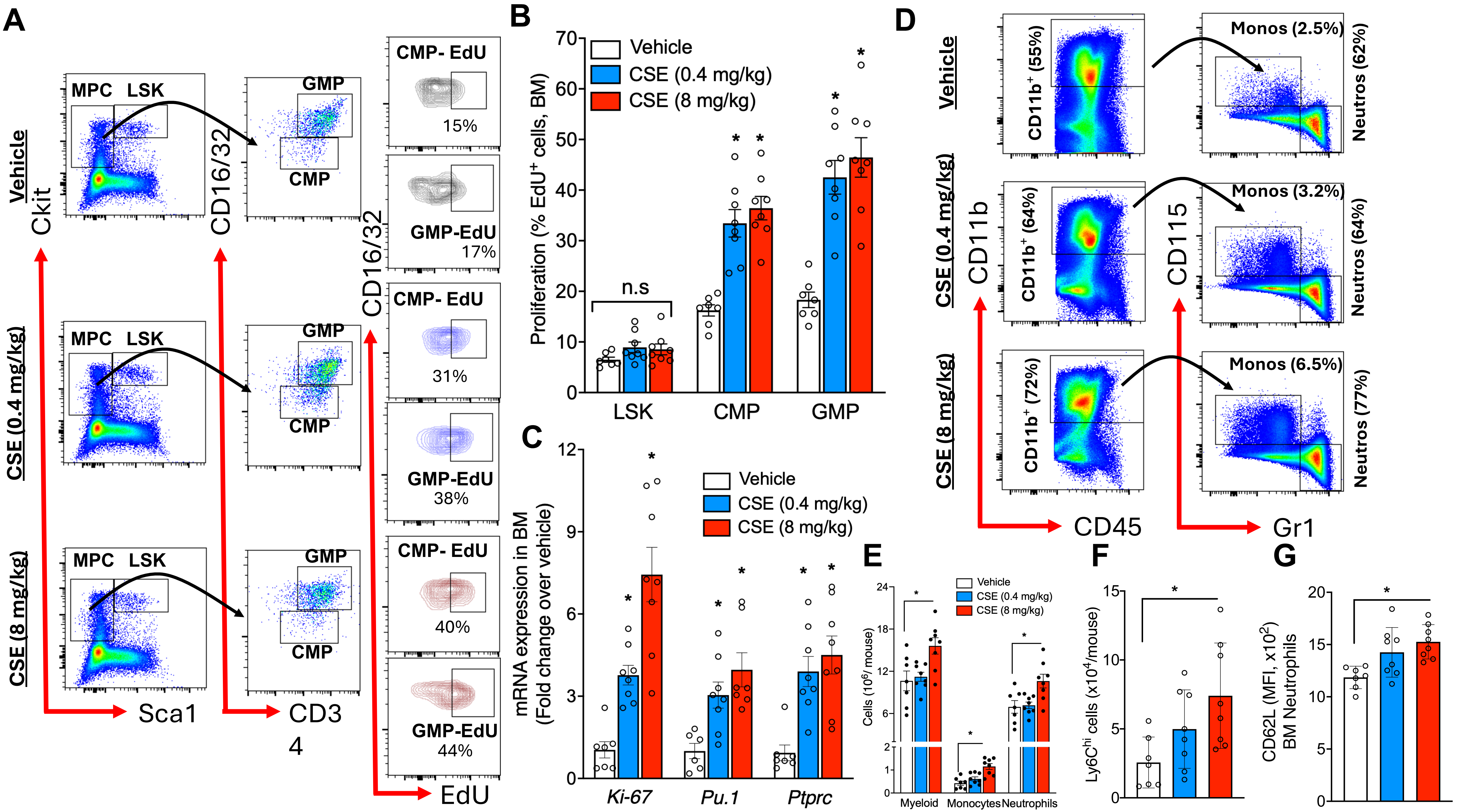

### Supplemental Figure-4

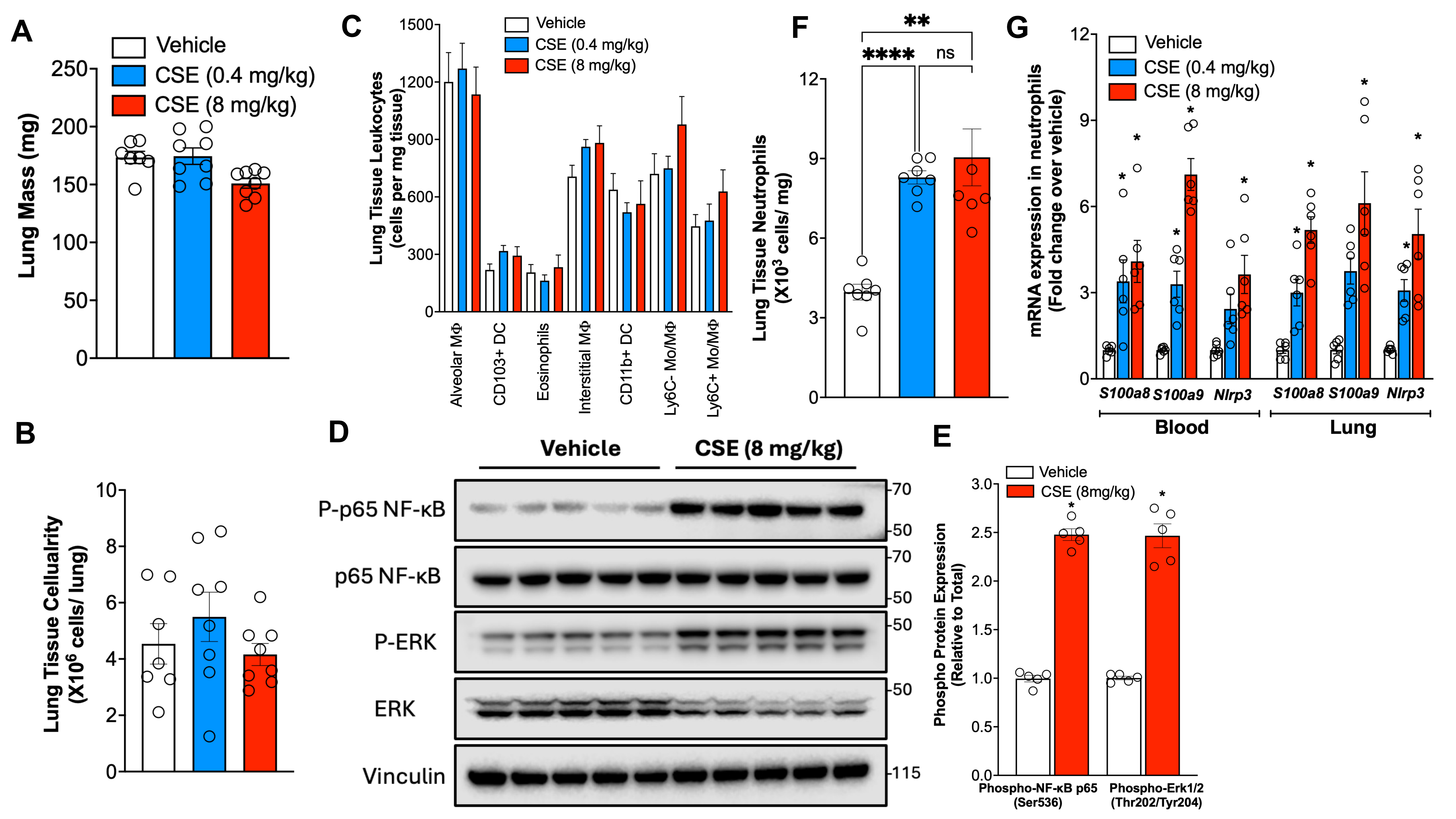

### Supplemental Figure-5

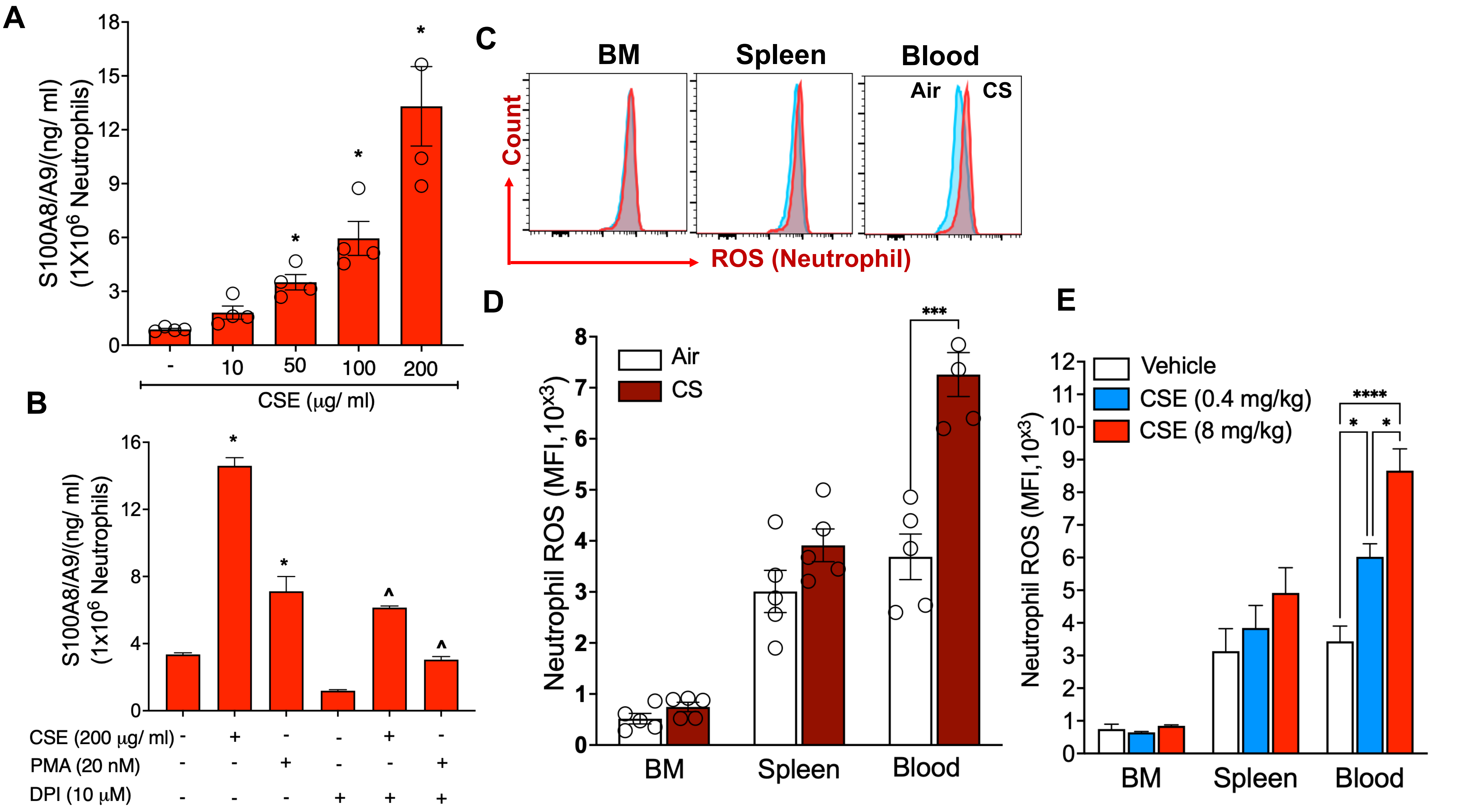

### Supplemental Figure-6

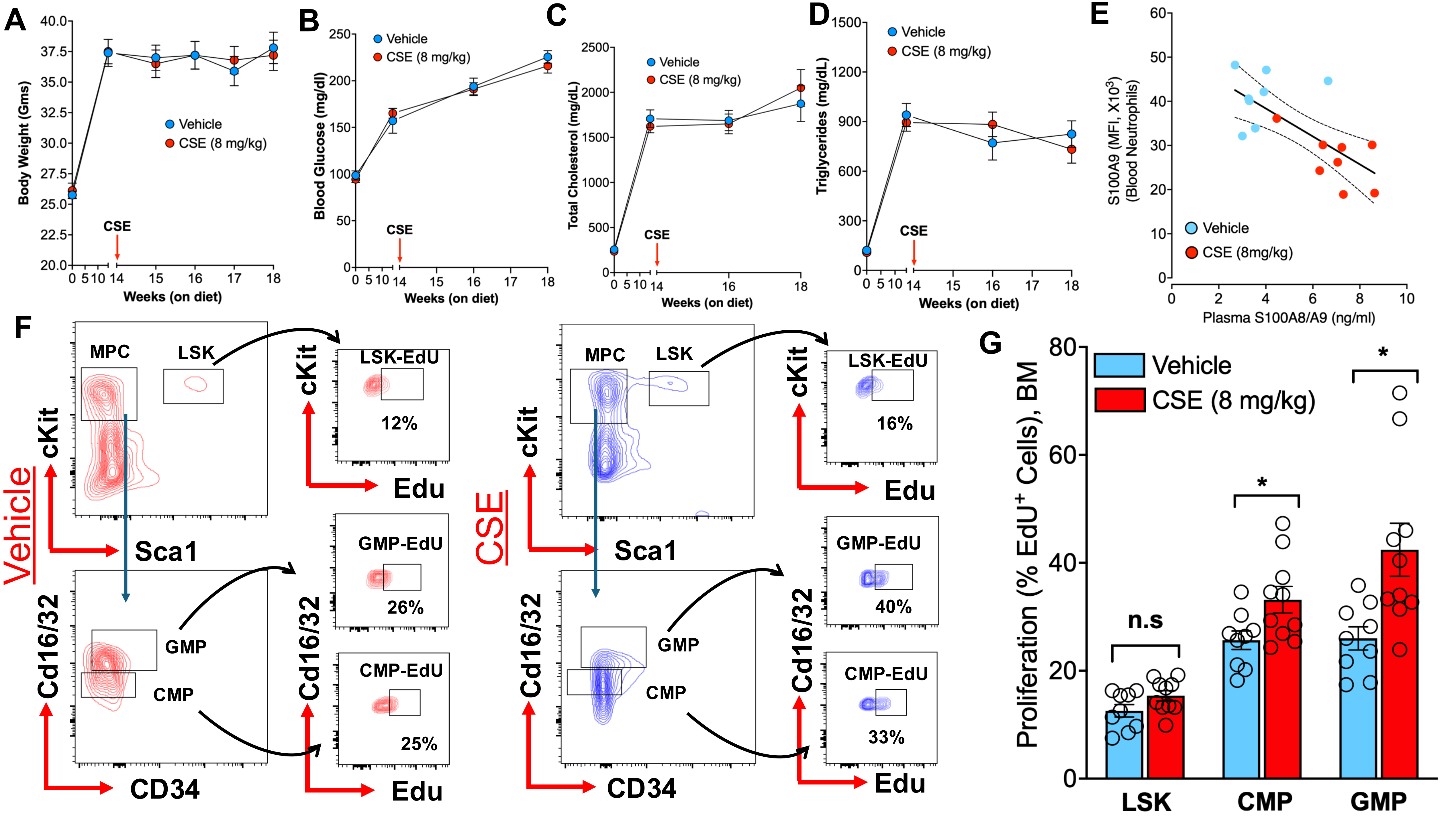

### Supplemental Figure-7

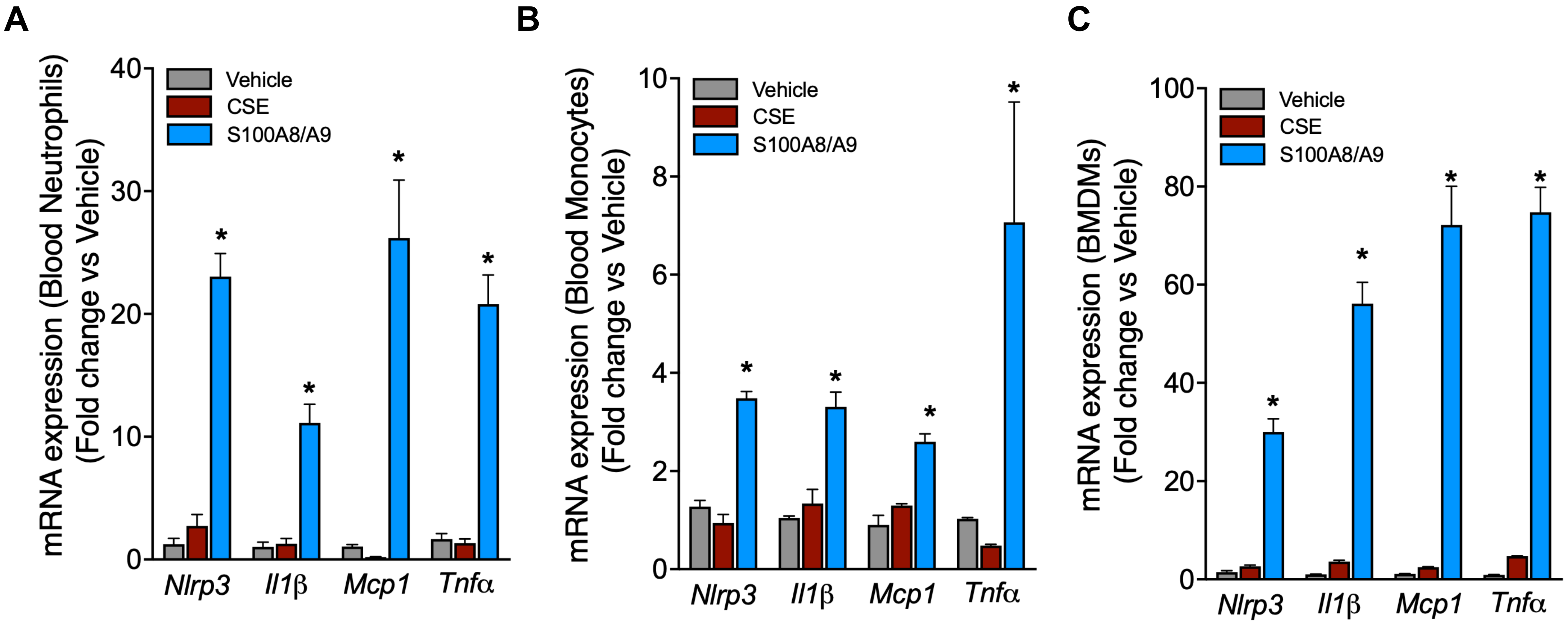

### Supplemental Figure-8

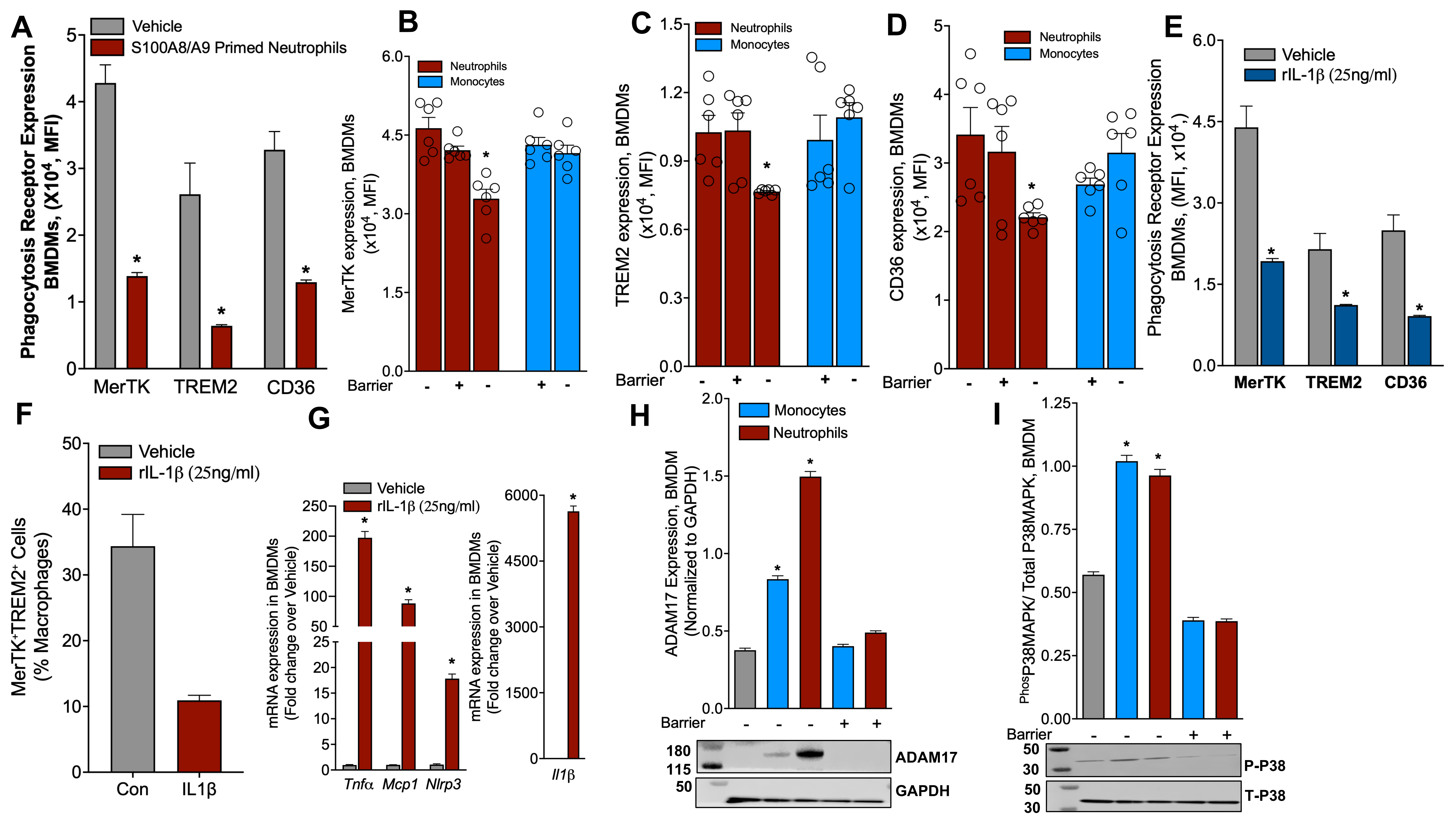

### Supplemental Figure-9

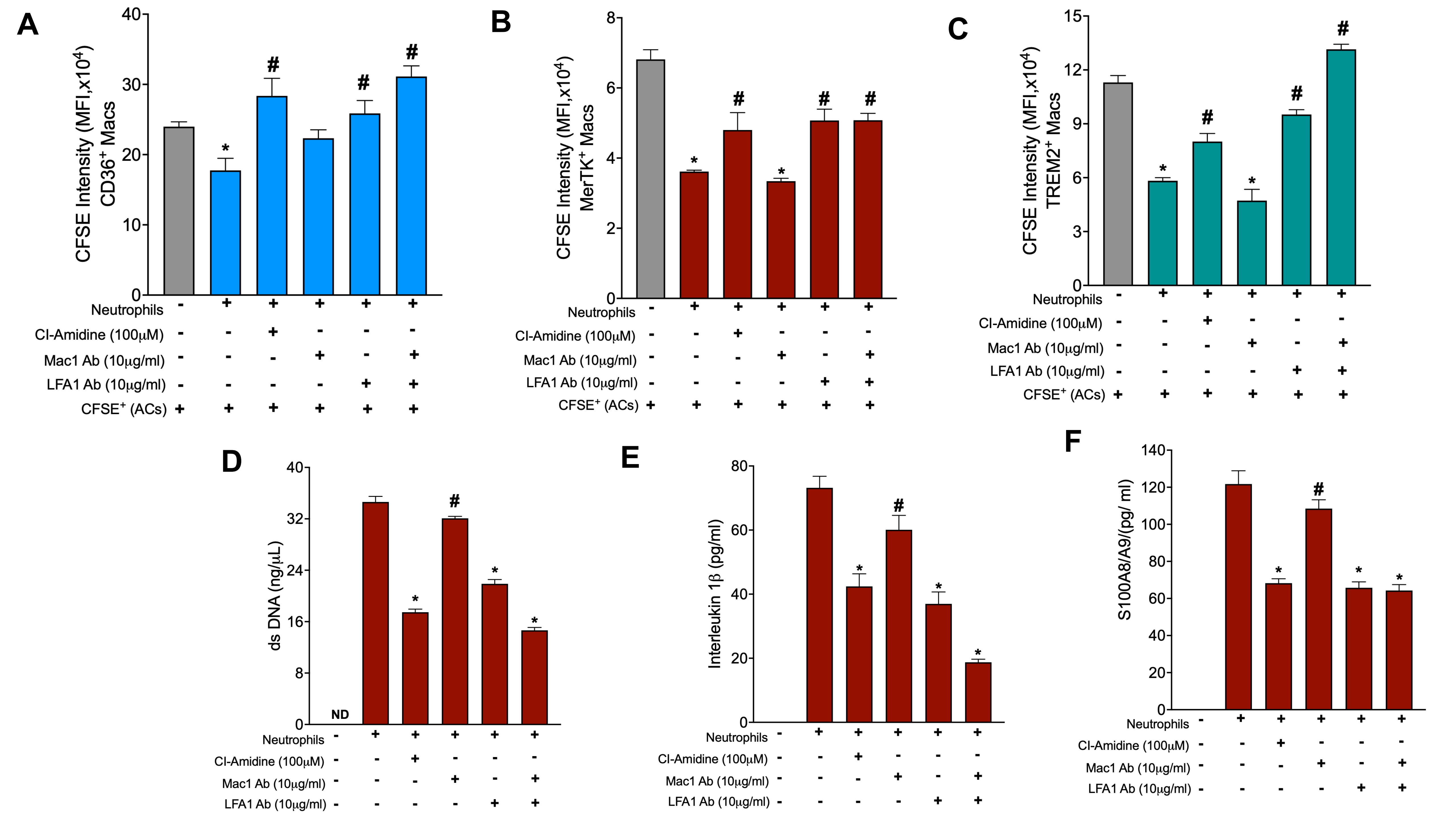

### Supplemental Figure-10

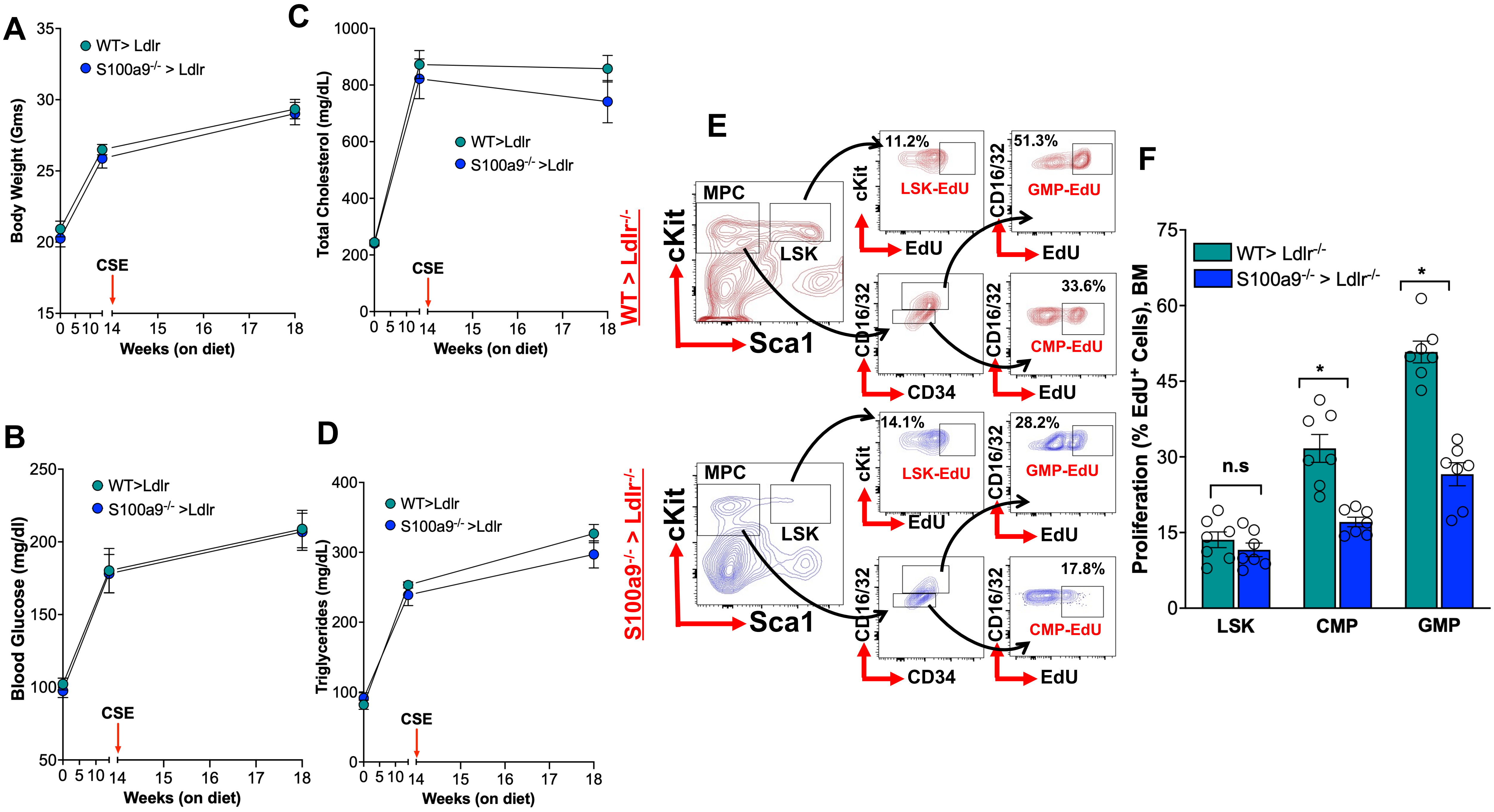
