## Supplemental Document for "Cigarette smoke aggravates atherosclerosis by promoting the infiltration of inflammasome-primed neutrophils and disrupting macrophage function in lesions"

**Short Title:** S100A8/A9 aggravate atherosclerosis

**Institutions**

[1]Department of Internal Medicine, Cardiovascular Section, University of Oklahoma Health Sciences Center, Oklahoma City, OK, USA

[2]Department of Surgery, Division of Cardiac Surgery, The Ohio State University Wexner Medical Center, Columbus, OH

[3] Baker Heart and Diabetes Institute, Division of Immunometabolism, Melbourne, Australia.

[^] These authors contributed equally

**Corresponding author,**

Dr. Prabha Nagareddy, PhD, FAHA

Department of Medicine, Cardiovascular Section

University of Oklahoma Health Sciences Center, Oklahoma City, USA

Ph: 495-271-8001 Ex 44737

ORCID: 0000-0002-0294-641X

**Mice and treatments**

Age (8-12 weeks) and sex matched Wild Type (C57BL/6J), *S100a9*$^{-/-}$ and *Ldlr*$^{-/-}$ mice were used for this study. All mice were maintained in pathogen-free environments of the University of Oklahoma HSC (OUHSC) / Ohio State University (OSU) animal facilities. Procedures were conducted under approval from the OUHSC and OSU's Institutional Animal Care and Use Committee, in accordance with the NIH Guide for the Care and Use of Laboratory Animals. All mice were fed with a normal chow diet or Wester type diet (WD) *ad libitum* and randomly assigned to experimental groups. All procedures were performed by personnel trained in the techniques according to IACUC guidelines. All invasive procedures were performed while animals were anesthetized. In compliance with the Animal Welfare Act principles, we reduced the number of animals sacrificed by omitting control groups for which comparative effects are already known.

**Method details**

**Exposure to Cigarette Smoke:** In this protocol, mice were exposed to side stream CS or air using whole-body exposure method. Mice were exposed chronically (4 or 8 weeks) to CS (50-200 µg/l total particulate matter) for up to 2 hours per day (1 hr. each time, twice), 5 days a week. At the end of exposure, a small volume of blood (0.2 ml) was drawn from tail artery to collect plasma and characterize the leukocyte profile. At the end of study, all mice were killed, and their blood, BAL fluids, spleen, BM and lungs were harvested for various biochemical measurements. Smoke-exposed mice were compared with air-exposed controls simultaneously.

**Cigarette Smoke Inhalation System:** A mixture of mainstream and side stream cigarette smoke was generated using type 3R4F research cigarettes (University of Kentucky, Lexington, KY) with a microprocessor-controlled TE10z smoking machine (Teague Enterprises, Woodland, CA) connected to whole-body exposure chambers. Mice were exposed to cigarette smoke for 4 hours per day (two 2-hour intervals), 6 days per week, for 4 weeks.

**Cigarette Smoke Extract In vivo studies:** Male, *Ldlr*$^{-/-}$ (B6.129S7-Ldlrtm1Her) mice on a C57BL/6 background were obtained from The Jackson Laboratory. Mice were housed in a temperature-controlled environment (25°C) under a 12-hour light/dark cycle. All mice were fed a high fat "Western" diet (Inotive TD.88137) for a total of 14 weeks. At the start of week 11, mice were randomly assigned to one of two treatment groups (n = 10 per group). The experimental group received cigarette smoke extract (CSE) at a dose of 8.0 mg/kg body weight, administered daily by oral gavage for four weeks. The control group received an equivalent volume of vehicle (phosphate-buffered saline, PBS) via the same route and schedule. All procedures were conducted in accordance with institutional guidelines and approved by the relevant animal care and use committee.

**Bone marrow transplant (BMT) studies**: Recipient mice were given 100mg/L neomycin 2 weeks before and after BMT. Briefly, *Ldlr*$^{-/-}$ mice were lethally irradiated (with 13 Gy from a Cesium gamma source in 2 doses separated by 4 hours) and transplanted with BM ($5x10^6$ cells) from donor mice. Chimerism was confirmed using either flow cytometry or qRT-PCR analysis of respective proteins/ genes (mRNA) in peripheral blood cells at ~6 weeks after transplantation. Once the chimerism was confirmed, mice were fed with Western type diet.

**White blood cell (WBC) counts**: Total WBC in freshly collected mouse blood was measured using hematology cell counter (Element HT-5, Heska Inc.,) or a K2 Fluorescent Viability Cell Counter (Nexcelom Biosciences).

**Flow cytometry**

**Blood and bone marrow leukocytes:** Leukocytes from the whole blood and BM were identified and quantified as described previously[1,2]. Briefly, fresh blood from mice was collected via tail bleeding directly into microcentrifuge tubes containing EDTA (5 mM) using heparin coated capillary tubes and immediately placed on ice. A small amount of blood was set aside for separation of plasma and for analysis of complete blood cell count (CBC) while the remaining blood was transferred to 15 ml centrifuge tubes and red blood cells were lysed using BD Pharm Lyse (BD Biosciences), washed, rinsed and resuspended in PBS or FACS buffer (Hanks balanced salt solution, HBSS + 0.1% BSA w/v, 5mM EDTA) for staining.

For isolation of BM, femurs and tibias from each mouse were isolated, cleaned and their ends were cut open with a sharp scissor to expose lumen. Bone marrow was flushed with ice cold PBS using a 23-gauge, 10 ml syringe directly through a 40 μM nylon mesh strainer on a 50 ml centrifuge tube. The BM samples were next spun at 300g for 10 min at 4ºC. Following aspiration of supernatant, red blood cells were lysed using RBC lysis buffer, washed twice with PBS and immediately stained with a fixable Live/Dead marker (1 μl per million cells in 1 ml for 30 minutes, Molecular Probes). The cells were then washed and resuspended in 100 μl of FACS buffer containing a cocktail of antibodies directed against different types of leukocytes. In experiments that required intracellular staining, the surface labelled cells were fixed and permeabilized with fix/ perm buffer (30 minutes) and then stained with intracellular antibodies (e.g. S100A9) in perm/ wash buffer. The stained cells were analyzed on a LSRII flow cytometer using FACS DiVa software. Once the doublets (by FSC-H vs. FSC-A) and dead cells were excluded, monocytes were identified as $CD45^+$ $CD115^+$ and further classified as Ly6-$C^{hi}$ and Ly6-$C^{lo}$; neutrophils as $CD45^+$ $CD115^-$ , $Ly6G^{hi}$; B lymphocytes as $CD45^+$, $CD11b^-$, $CD19^+$ and T lymphocytes as $CD45^+$, $CD11b^-$, $CD3^+$, $CD4^+$ or $CD8^+$ cells.

**Spleen leukocytes:** The spleens were weighed and cut into small pieces, mashed using the rear end of a 1 ml syringe piston and passed through 40 μM nylon mesh strainer and the suspension was centrifuged at 10 minutes at 300 x g, 4°C. The resulting single cell suspensions were rinsed, resuspended in RBC lysis buffer. After RBC lysis and cell count, the cells were stained for viability, surface and intracellular proteins as described above. Neutrophils were identified as $CD45^+$, $Ly6G^{hi}$ cells and monocytes as $CD45^+$, $Ly6G^-$, $F4/80^-$, $CD11b^+$, $CD115^+$ and further classified as $Ly6\text{-}C^{hi}$ and $Ly6\text{-}C^{lo}$ cells.

**Hematopoietic stem cells in the BM and spleen:** Hematopoietic stem and progenitor cells (HSPCs) from the BM and spleen were prepared as described above. After cell count and staining for Live/Dead markers, the cells were resuspended in a cocktail of antibodies first against lineage-committed cells (B220, CD19, CD11b, CD3e, TER-119, CD2, CD8, CD4, Ly6-C/G) followed by markers of stem cells, proliferation (e.g. Ki67) and EdU. Hematopoietic stem and progenitor cells were first identified as $Lin^-$, $Sca1^+$ and $ckit^+$ (LSK) while the hematopoietic progenitor subsets were separated by using antibodies to CD16/CD32 (FcγRII/III) and CD34. Common Myeloid Progenitors (CMP) were identified as $Lin^-$, $Sca1^-$, $ckit^+$, $CD34^{int}$, $Fc\gamma RII/III^{int}$, Granulocyte Macrophage Progenitors (GMP) as $Lin^-$, $Sca1^-$, $ckit^+$, $CD34^{int}$, $Fc\gamma RII/III^{hi}$. Common lymphoid progenitors (CLP) were identified as $Lin^-$, $Sca1^{lo}$, $ckit^{lo}$ and IL7R+ cells. Hematopoietic stem cells (HSCs) were identified as $Lin^-$, $Sca1^+$, $ckit^+$, $CD48^-$ and $CD150^+$. Cellular proliferation in hematopoietic stem and progenitor cells was assessed by measuring the incorporation of EdU and /or Ki67 according to the manufacturer's protocol (Click-iT™ Plus EdU, Molecular Probes).

**In vivo proliferation assay (EdU)**: For proliferation studies, mice were injected with 0.2 mg of 5-(ethynyl-2´-deoxyuridine) EdU ~12-14 Hrs. *via* tail vein prior to sacrifice. In preparation for flow cytometry, cell populations were immunostained as described above and the incorporation of EdU was quantified using Click-iT™ Plus EdU flow cytometry assay kit (Molecular Probes, Eugene OR) according to the manufacturer's instructions. Proliferation was quantified and expressed as percentage of $EdU^+$ cells.

**Ex vivo proliferation assay (EdU):** Fresh BM cells from WT mice were isolated and resuspended in IMDM (Invitrogen) containing 10% FCS (StemCell Technologies) and cultured for 2 Hrs in tissue culture flasks to enrich stem and progenitor cells (non-adherent cells). Following the removal of adherent cells, the remaining cell were cultured in IMDM containing stem cell factor (100ng/ml), IL-3 (6ng/ml) and GM-CSF (2ng/ml) for 12-16 hours. For proliferation measurements, cells were treated with S100A8/A9 (2 µg/ml) and varying concentrations of CSE for 16 hrs. in the presence of 10 µM EdU. HSCs, HSPCs and neutrophils in the culture were determined by flow cytometry. Proliferation was measured by determining the amount of EdU in each cell type and represented as % $EdU^+$ cells.

**Plasma biochemistry and cytokine Levels**: Plasma samples were obtained from 6-h–fasted mice. Blood glucose was measured directly from the tail tip using a One Touch Ultra 2 glucose monitoring system (Lifescan, Johnson & Johnson). Total cholesterol (TC) and triglycerides (TG) levels in the plasma were measured using colorimetric assays (Wako Diagnostics). Plasma S100A8/A9 was measured using the mouse S100A8/S100A9 Heterodimer DuoSet ELISA (R&D Systems)

**Assessment of atherosclerosis:** For atherosclerotic lesion analysis, serial frozen sections of proximal aortic roots were prepared and stained with H&E for quantification of lesion size as previously described[3]. Oil Red O staining was employed for lipid quantification while CD68 staining was used to assess macrophages in lesions of the proximal aortic root. *En face* analysis was performed to quantify lipids in atherosclerotic plaques of aortic arch. Quantification was performed using either Image J, Image Pro Plus or Adobe Photoshop CS5.

**Assessment of leukocytes in aortic root lesions:** Mouse aortas were dissected following saline perfusion to remove circulating blood. The aortas were gently cleaned to remove excess fat and connective tissue, then placed in 970 µL of RPMI medium containing 0.03 mg/mL Liberase (Sigma, Cat# 5401119001) in a 2 mL Eppendorf tube. The tissue was finely minced using sharp scissors and incubated at 37°C for 45 minutes with gentle shaking at 100 rpm to allow enzymatic digestion. Following digestion, the cell suspension was filtered through a 100 µm cell strainer into a 50 mL Falcon tube, and residual tissue was rinsed through the strainer using FACS buffer. The filtered suspension was centrifuged at 500 × g for 5 minutes, and the supernatant was discarded. The cell pellet was resuspended in 1 mL of wash buffer, spun again at 500 × g for 5 minutes, and the final pellet was resuspended for downstream flow cytometry staining.

**Neutrophil-Macrophage**

**Co-culture Studies:** Bone marrow-derived neutrophils and monocytes were isolated from Wild-type C57BL/6 mice and stimulated with LPS (1 µg/mL) or vehicle for 3 hours. Following stimulation, cells were washed twice with PBS to remove residual LPS. For indirect co-culture, LPS-primed neutrophils ($1×10^6$ cells) or monocytes ($0.8×10^6$ cells) were seeded into 0.4 µm pore size cell culture inserts (Sigma, Cat# CLS3450) and placed into 6-well plates containing bone marrow–derived macrophages (BMDMs; $2×10^6$ cells). Cells were co-cultured overnight. In parallel, for direct co-culture, the same number of LPS-primed neutrophils or monocytes were added directly to BMDMs and cultured under identical conditions. After overnight incubation, BMDMs were washed thoroughly with PBS to remove non-adherent cells, scraped from the culture wells, and processed for flow cytometric analysis.

**RNA isolation, cDNA synthesis and qRT-PCR :** Total RNA from cells was extracted using either RNeasy Micro kit (Qiagen) or PureLink RNA mini kit (Ambion). RNA quality and quantity were assessed on a DS-11 FX nanoscale spectrophotometer (Denovix Inc.) or 2100 Bioanalyzer (Agilent Technologies Inc., Santa Clara CA). cDNA was synthesized using SuperScript VILO cDNA synthesis kit from Invitrogen (ThermoFisher Scientific). qRT-PCR was monitored in real time with a Quant Studio Real-Time PCR system using SYBR Green Reagents (Life Technologies). mRNA levels were normalized to either 18S or GAPDH and data computed using using the $2^{-DDCT}$ method. The sequences for primers used in qRT-PCR are listed in key resources table.

**Statistics:** Normality of data distribution was assessed using the Shapiro–Wilk test. For comparisons between two groups, an unpaired two-tailed t-test was used for normally distributed data, whereas a Mann–Whitney test was applied for non-normally distributed variables. For comparisons involving three or more groups with normally distributed data, one-way or two-way analysis of variance (ANOVA) was performed, followed by Holm–Šidák or Tukey's multiple-comparison tests as appropriate. For nonparametric datasets, the Kruskal–Wallis test followed by Dunn's multiple-comparison test was used. Correlations and linear relationships between variables were analyzed using Spearman's rank correlation, and the goodness of fit was expressed as the coefficient of determination ($R^2$). A p value $\leq 0.05$ was considered statistically significant throughout all analyses. All statistical analyses were performed using GraphPad Prism version 10 (GraphPad Software, San Diego, CA).

**Legends to supplemental figures.**

**Figure S1. Effects of cigarette smoke (CS) exposure on splenic hematopoiesis and neutrophil alarmin expression. A)** Quantification of hematopoietic stem and progenitor cells (HSPCs) and their **B**) proliferation (as determined by EdU incorporation) in spleens of air- and CS-exposed mice using flow cytometry. C) Quantification of splenic monocytes and neutrophils. **D**) Measurement of spleen weight at termination following 4 weeks of CS exposure. **E**) Intracellular levels of S100A8/A9 in splenic neutrophils as assessed by flow cytometry. Data represent mean ± SEM. Statistical analysis for panels A-E was performed using unpaired two-tailed *t*-test (*$p < 0.05$ vs. Air).

**Figure S2. Cigarette smoke (CS) exposure induces inflammatory remodeling of the lung. A**) Representative flow cytometry plots and gating strategy used to identify and quantify leukocyte populations in lung tissue at termination. All flow cytometry data were gated to exclude debris, dead cells, and doublets. The numbers in parentheses indicate the percentage of each identified population relative to its parent population. Quantification of neutrophils (**B**) and other leukocyte subsets (**C**), including interstitial macrophages (IMs), alveolar macrophages (AMs), dendritic cell (DC) subsets ($CD103^+$ and $CD11b^+$), eosinophils, and $Ly6C^+$/$Ly6C^-$ monocytes in lung tissue from air- and CS-exposed mice. D) Quantification of leukocyte populations in bronchoalveolar lavage fluid (BALF), showing increased total leukocytes, neutrophils, $CD103^+$ DCs, IMs, and $Ly6C^+$ monocytes following CS exposure. **E**) Quantification of S100A8/A9 protein levels in BALF by ELISA. **F**) mRNA expression of inflammatory markers in lung tissue. Results are expressed as mean ± SEM. Statistical analysis for panels B-F was performed unpaired two-tailed *t*-test (*$p < 0.05$ vs. Air).

**Figure S3. Oral cigarette smoke extract (CSE) administration enhances bone marrow (BM) myelopoiesis in a dose-dependent manner**. **A**) Representative flow cytometry plots showing multipotential progenitor cells (lineage$^-$, c-Kit$^+$, and Sca-1$^+$; LSK), common myeloid progenitors (CMPs), and granulocyte–macrophage progenitors (GMPs) and their **B**) quantification (as assessed by EdU incorporation) in BM of control and CSE-treated mice. All flow cytometry data were gated to exclude debris, dead cells, and doublets. **C**) Expression of proliferation and myeloid lineage markers, including Ki-67, Pu.1, and Ptprc, in BM cells. **D**) Representative flow cytometry plots and E) quantification of total myeloid cells (CD45$^+$, CD11b$^+$), monocytes, and neutrophils in BM following CSE exposure. F) Quantification of Ly6C$^{hi}$ proinflammatory monocytes and **G**) Cell surface expression of CD62L as assessed by mean fluorescence intensity (MFI) using flow cytometry. Results are presented as mean ± SEM. Statistical analysis for panels B, C, E, F, and G was performed using an unpaired two-tailed *t*-test (**p* < 0.05 vs. vehicle; n.s., not significant).

**Figure S4. Oral cigarette smoke extract (CSE) exposure induces mild lung inflammation without altering lung mass or immune cell composition**. **A**) Measurement of lung mass in control and CSE-treated mice at termination. **B**) Total lung cellularity determined by flow cytometry. **C**) Quantification of immune cell subsets in lung tissue, including alveolar macrophages, interstitial macrophages, dendritic cells, monocytes, and neutrophils. **D**), Representative immunoblots of lung tissue showing phosphorylation of p65–NFκB and ERK (indicators of cellular activation) and corresponding **E**) quantification, demonstrating mild inflammatory changes following oral CSE administration. **F**) Flow cytometric quantification of neutrophils infiltrating lung tissue. **G**) Comparative analysis of inflammatory marker expression (mRNA) in neutrophils isolated from blood and lung of CSE-treated mice. Results are presented as mean ± SEM. Statistical analysis for panels **A, B, C** and **G** was performed using one-way ANOVA and Tukey's multiple comparison test (*p < 0.05, vs. vehicle). For **F**, one-way ANOVA with Dunnett's T3 multiple comparison test (**** p< 0.0001, ** p<0.001 vs Vehicle, n.s., not significant). For **E**, unpaired two-tailed *t*-test (*p* < 0.05 vs. vehicle in the corresponding protein).

**Figure S5. Cigarette smoke extract (CSE) induces oxidative stress–mediated release of S100A8/A9 from neutrophils. A**) Quantification of S100A8/A9 secretion from bone marrow neutrophils treated with increasing concentrations of CSE. **B**) Comparison of S100A8/A9 release following treatment with CSE or phorbol myristate acetate (PMA) in the presence or absence of diphenyleneiodonium (DPI), an NADPH oxidase inhibitor. **C**) Histogram representing the mean fluorescence intensity(MFI) of reactive oxygen species (ROS) and its **D**) quantification in blood, BM, and splenic neutrophils of air- and cigarette smoke (CS)- exposed mice. **E**) ROS levels as assessed by MFI in neutrophils from blood, BM, and spleen of vehicle- and CSE-treated mice. Results are presented as mean ± SEM. Statistical analysis for panels **A**, **B** and **E** was performed using one-way ANOVA and Tukey's multiple comparison test (*p < 0.05, vs. vehicle, **** p< 0.0001, ^p< 0.05 vs CSE along group). For **D**, unpaired two-tailed *t*-test (****p* < 0.005 vs Air).

**Figure S6. Oral cigarette smoke extract (CSE) administration enhances myelopoiesis without affecting metabolic parameters in atherosclerotic mice**. Body weight (**A**), blood glucose (**B**), total cholesterol (TC; **C**), and triglyceride (TG; **D**) levels in $Ldlr^{-/-}$ mice fed a Western diet (WD) for 14 weeks, followed by daily treatment with vehicle or high-dose CSE (8 mg/kg) for an additional 4 weeks. **E**) Linear regression analysis showing the relationship between plasma S100A8/A9 levels (X-axis) and neutrophil S100A9 mean fluorescence intensity (MFI) (Y-axis). Each point represents an individual mouse, with blue dots indicating vehicle group while red dots indicating CSE group. A linear regression line is included for visualization. **F**) Representative flow cytometric plots and gating strategy used to identify, and **G**) quantify, the proliferation of common myeloid progenitors (CMPs), granulocyte-macrophage progenitors (GMPs) and hematopoietic stem and progenitor cell (LSKs) in response to CSE treatment. All flow cytometry data were gated to exclude debris, dead cells, and doublets. The numbers in parentheses indicate the percentage of EdU+ cells within each population relative to its parent population. Results are presented as mean ± SEM. Statistical analysis for panel **E** was performed using Spearman's rank correlation.

Plasma S100A8/A9 levels were inversely correlated with neutrophil S100A9 mean fluorescence intensity (MFI) across all samples ($R^2 = 0.464$, slope = −3.38, $p = 0.015$), indicating that higher circulating S100A8/A9 is associated with reduced neutrophil S100A9 expression. For panel **G**, comparisons were made using an unpaired two-tailed t-test (*$p < 0.05$ vs. vehicle, n.s, not significant).

**Figure S7. S100A8/A9, but not cigarette smoke extract (CSE), activates inflammasome signaling in myeloid cells**. Gene expression analysis of inflammasome and inflammatory markers (*Nlrp3, Il1β, Tnfα,* and *Mcp1*) in sorted blood monocytes (**A**) and neutrophils (**B**) following exposure to CSE or recombinant S100A8/A9. **C**) mRNA expression of the inflammasome and inflammatory markers in bone marrow–derived macrophages (BMDMs) treated with CSE or S100A8/A9. Results are presented as mean ± SEM. Statistical analysis for panels **A** to **C** was performed using one-way ANOVA and Tukey's multiple comparison test (*$p < 0.05$, vs. all other groups for each gene).

**Figure S8. IL-1β and S100A8/A9 released from neutrophils impair macrophage phagocytic receptor expression through p38 MAPK–ADAM17 signaling.** (**A**) Expression of phagocytic receptors (PRs: MerTK, TREM2, and CD36) in bone marrow–derived macrophages (BMDMs) co-cultured with S100A8/A9-primed blood neutrophils. Expression of MerTK (**B**), TREM2 (**C**), and CD36 (**D**) assessed by mean fluorescence intensity (MFI) using flow cytometry in BMDMs following co-culture with inflammasome-primed monocytes or neutrophils. Stimulation of BMDMs with recombinant IL-1β showing downregulation of PRs (**E**), reduction in functional $MerTK^+TREM2^+$ macrophages (**F**), and acquisition of a pro-inflammatory phenotype (**G**) determined by mRNA expression of inflammatory markers. Immunoblot analysis of ADAM17 protein (**H**) and phosphorylated p38 MAPK (**I**) in BMDMs co-cultured with inflammasome-primed neutrophils or monocytes, indicating activation of the IL-1β–p38 MAPK-ADAM17 signaling axis.

Results are presented as mean ± SEM. Statistical analysis for panels **A, E, F**, and **G** was performed using an unpaired two-tailed t-test (*$p < 0.05$ vs. vehicle). Statistical analysis for panels **B, C**, and **D** was performed using one-way ANOVA followed by Tukey's multiple-comparison test (*$p < 0.05$ vs. neutrophils with and without barrier; first and second maroon bars). Statistical analysis for panels **H** and **I** was performed using one-way ANOVA followed by Tukey's multiple-comparison test ($p < 0.05$ vs. control; first gray bar).

**Figure S9. $\beta_2$ integrin blockade or NETosis inhibition preserves macrophage phagocytic receptor expression and efferocytosis function**. Quantification of efferocytosis capacity in macrophage subpopulations expressing high levels of CD36 (**A**), MerTK (**B**), or TREM2 (**C**), as assessed by the mean fluorescence intensity (MFI) of internalized CFSE-labeled apoptotic cells. Quantification of NETosis (**E**), IL-1β release (**F**), and S100A8/A9 secretion (**G**) under indicated treatment conditions. Results are presented as mean ± SEM. Statistical analyses for panels **A–C** were performed using one-way ANOVA followed by Tukey's multiple-comparison test (*$p < 0.05$ vs. neutrophils alone, first gray bar; $^{\#}p < 0.05$ vs. neutrophils + CFSE-labeled apoptotic cells, second bar). For panels **D–E**, analyses were conducted using one-way ANOVA followed by Tukey's multiple-comparison test (*$p < 0.05$ vs. neutrophils alone, second bar; $^{\#}p < 0.05$ vs. neutrophils + Cl-Amidine, third bar).

**Figure S10. Deletion of S100a9 attenuates CSE-induced myelopoiesis without affecting systemic metabolic parameters**. Measurement of body weight (**A**), serum glucose (**B**), total cholesterol (**C**), and low-density lipoprotein (LDL) cholesterol (**D**) in $Ldlr^{-/-}$ recipient mice reconstituted with wild-type (WT) or $S100a9^{-/-}$ bone marrow (BM), fed a Western diet (WD) for 14 weeks, and treated with cigarette smoke extract (CSE) for an additional 4 weeks. **E**, Representative flow cytometry plots showing the gating strategy used to identify hematopoietic stem and progenitor cells (LSKs) and their **F**) proliferation as assessed by EdU incorporation.

LSKs were defined as Lineage$^{-}$, c-Kit$^{+}$, and Sca-1$^{+}$ cells; CMPs (common myeloid progenitors) and GMPs (granulocyte–macrophage progenitors) were identified based on standard surface markers. All flow cytometry data were gated to exclude debris, dead cells, and doublets. The numbers in parentheses indicate the percentage of EdU$^{+}$ cells relative to their parent population. Results are presented as mean ± SEM. Statistical analyses for panels **A–D** were performed using two-way ANOVA, followed by Bonferroni post hoc test. For panel **F**, an unpaired two-tailed t-test was used ($p < 0.05$ vs. S100a9$^{-/-}$ BMT group, n.s, not significant).

**Supplemental Table S1: Key Resources and Reagents**

| REAGENT or RESOURCE | SOURCE | IDENTIFIER |
|---|---|---|
| **Antibodies** | | |
| Anti-mouse CD45 (clone 30-F11) | BioLegend | 103116 |
| Anti-mouse CD11b (clone M1/70) | BioLegend | 101228 |
| Anti-mouse CD115 (clone AFS98) | BioLegend | 135515 |
| Anti-mouse Ly6-G (clone 1A8) | BioLegend | 127612 |
| Anti-mouse Ly6-C (clone HK1.4) | BioLegend | 128018 |
| Anti-mouse Gr1 (clone RB6-8C5) | ThermoFisher | 108428 |
| Anti-mouse B220 (clone RA3-6B2) | ThermoFisher | 11-0452-85 |
| Anti-mouse CD11b (clone M1/70) | BD Biosciences | 553310 |
| Anti-mouse CD19 (clone 1D3/CD19) | BioLegend | 152404 |
| Anti-mouse CD3e (clone 145-2C11) | ThermoFisher | 11-0031-85 |
| Anti-mouse TER-119 (clone TER-119) | ThermoFisher | 11-5921-85 |
| Anti-mouse CD2 (clone RM2-5) | ThermoFisher | 11-0021-85 |
| Anti-mouse CD8a (clone 53-6.7) | ThermoFisher | 11-0081-85 |
| Anti-mouse CD4 (clone GK1.5) | ThermoFisher | 11-0041-85 |
| Anti-mouse Gr1 (clone RB6-8C5) | ThermoFisher | 11-5931-85 |
| Anti-mouse CD150 (clone TC15-12F12.2) | BioLegend | 115922 |
| Anti-mouse CD34 (clone 581) | BioLegend | 343512 |
| Anti-mouse cKit (clone 2B8) | BioLegend | 105835 |
| Anti-mouse CD16/32 (clone 93) | BioLegend | 101308 |
| Anti-mouse Sca1 (clone D7) | BioLegend | 108114 |
| Anti-mouse CD48 (clone HM48-1) | BioLegend | 103432 |
| Anti-mouse CD45.2 (clone 104) | ThermoFisher | 25-0454-82 |

| | | |
|---|---|---|
| MERTK Super Bright 436 (DS5MMER) | ThermoFisher | 62-5751-82 |
| Anti-mouse CD45.1 (clone A20) | ThermoFisher | 12-0453-82 |
| Human/MouseTREM2 APC-conjugated Antibody | R&D Systems | FAB17291A |
| Anti-mouse Annexin V | BioLegend | 640943 |
| Anti-CD68 antibody produced in rabbit | Sigma-Aldrich | SAB5700832 |
| Anti-mouse Ki67 (clone SolA15) | ThermoFisher | 12-5698-82 |
| **Chemicals, Peptides, and Recombinant Proteins** | | |
| Fixable Live/Dead marker | ThermoFisher | C2926 |
| LPS from E. coli | Enzo Life Sciences | ALX-581-012 |
| Acridine orange & Propidium Iodide) | Nexcelom | tlrl-vad |
| Weigert's iron hematoxylin solution A | EMS | 26758-01 |
| Weigert's iron hematoxylin solution B | EMS | 26758-02 |
| Collagenase I | Sigma-Aldrich | C0130 |
| Collagenase XI | Sigma-Aldrich | C7657 |
| DNAse I | Sigma-Aldrich | D5319 |
| Hyaluronidase | Sigma-Aldrich | H3506 |
| DAPI | ThermoFisher | F10347 |
| TNF-α | Sigma-Aldrich | T7539 |
| Liberase TM | Sigma-Aldrich | 540111901 |
| **Critical Commercial Assays** | | |
| Mouse Neutrophil Isolation kit | Miltenyi Biotec | 130-097-658 |

| | | |
|---|---|---|
| BD Pharm Lyse | BD Biosciences | 555899 |
| Click-iT™ Plus EdU Kit | ThermoFisher | C10640 |
| Ambion Purelink RNA Mini Kit | ThermoFisher | 12183025 |
| SuperScript IV VILO Master Mix | ThermoFisher | 11766050 |
| TaqMan Fast Advanced Master mix | ThermoFisher | 4444963 |
| Fast SYBR Green Master Mix | ThermoFisher | 4385612 |
| Total RNA Isolation Kit | ThermoFisher | AM1931 |
| Mouse IL-1 beta Uncoated ELISA Kit | ThermoFisher | 88-7013A-88 |
| Mouse S100A8/A9 Heterodimer DuoSet Elisa | R&D Systems | DY8596-05 |
| Mouse Cytokine Array Q1 | RayBiotech | QAM-CYT-1-1 |
| QuantiFlour One dsDNA System | Promega | E4871 |
| BCA Protein Assay kit | ThermoFisher | 23225 |
| **Animals** | | |
| C57BL/6J | JAX Laboratories | 000664 |
| B6.129S7-*Ldlr*$^{tm1Her}$/J (Ldlr KO) | JAX Laboratories | 002207 |
| **Software** | | |
| FACS DiVa | BD Biosciences | http://www.bdbiosciences.com |
| FlowJo software (Ashland, OR) | FlowJo | http://www.flowjo.com |
| NIH ImageJ | NIH | http://imagej.net |
| Gen5 | BioTek | https://www.biotek.com |

| | | |
|---|---|---|
| Spectraflo | Cytek | https://cytekbio.com/ |
| **Equipment/ Instruments** | | |
| Vevo 3100 High-resolution US imaging system | VisualSonics Inc., | https://www.visualsonics.com |
| K2 Fluorescent Viability Cell Counter | Nexcelom | https://www.nexcelom.com |
| LSRFortessa | BD Biosciences | |
| LSRII Flow Cytometer | BD Biosciences | https://www.bdbiosciences.com/en-in/instruments/research-instruments/research-cell-analyzers/lsrfortessa |
| Cytek Northern Lights | Cytek | https://cytekbio.com/pages/northern-lights |
| GentleMACS Octo Dissociator | Miltenyi Biotec | https://www.miltenyibiotec.com |
| ECLIPSE Ti2 | Nikon | https://www.microscope.healthcare.nikon.com/products/inverted-microscopes/eclipse-ti2-series |
| Digital Slide Scanner Axioscan | Zeiss | https://www.zeiss.com/microscopy/us/products/imaging-systems/axioscan-7.html |

| | | |
|---|---|---|
| Element HT5 | Heska Inc., | https://www.heska.com/product/element-ht5/ |
| Micro- Cell | MicroVet Diagnostics | https://www.microvetdiagnostics.com/Products/Hematology |

**Supplemental Table S2: Taqman gene expression assays for qPCR analysis**

| No. | Gene | Assay ID | Catalog No. | Vendor |
|---|---|---|---|---|
| 1 | IL-1B | Mm00434228_m1 | 4331182 | Applied Biosystems |
| 2 | TNF-A | Mm00443258_m1 | 4331182 | Applied Biosystems |
| 3 | MCP-1 | Mm00441242_m1 | 4331182 | Applied Biosystems |
| 4 | Nlrp3 | Mm00840904_m1 | 4331182 | Applied Biosystems |
| 5 | Eukaryotic 18S rRNA Endogenous Control | | 4319413E | Applied Biosystems |

### References


1. Nagareddy PR, Kraakman M, Masters SL, Stirzaker RA, Gorman DJ, Grant RW, Dragoljevic D, Hong ES, Abdel-Latif A, Smyth SS, et al. Adipose tissue macrophages promote myelopoiesis and monocytosis in obesity. *Cell Metab*. 2014;19:821-835. doi: 10.1016/j.cmet.2014.03.029
2. Nagareddy PR, Murphy AJ, Stirzaker RA, Hu Y, Yu S, Miller RG, Ramkhelawon B, Distel E, Westerterp M, Huang LS, et al. Hyperglycemia promotes myelopoiesis and impairs the resolution of atherosclerosis. *Cell Metab*. 2013;17:695-708. doi: 10.1016/j.cmet.2013.04.001
3. Al-Sharea A, Murphy AJ, Huggins LA, Hu Y, Goldberg IJ, Nagareddy PR. SGLT2 inhibition reduces atherosclerosis by enhancing lipoprotein clearance in Ldlr(-/-) type 1 diabetic mice. *Atherosclerosis*. 2018;271:166-176. doi: 10.1016/j.atherosclerosis.2018.02.028
